# The microbiota impacts life history traits and mating success in male *Aedes aegypti* mosquitoes

**DOI:** 10.64898/2026.03.09.710597

**Authors:** Noha K. El-Dougdoug, Dom Magistrado, Kayla I. Perry, Sarah M. Short

## Abstract

*Aedes aegypti* mosquitoes transmit multiple arboviruses, but population suppression through the mass release of sterile or incompatible male mosquitoes can effectively reduce populations. These methods depend on the reliable mass-rearing of healthy, robust males that can successfully mate with wild females. The microbiota, a critical component of the larval diet, can dramatically influence life history traits relevant to mass-rearing and male quality. Here, we used axenic (microbe-free), monoxenic (inoculated with *E. coli*), and “laboratory community” mosquitoes (inoculated with an undefined microbiota derived from laboratory mosquitoes) to show that longevity was significantly enhanced in axenic and monoxenic males compared to laboratory community males. Moreover, monoxenic males more efficiently obtained mates in non-competitive mating scenarios compared to laboratory community males. However, microbiota treatment had no effect when males from different treatments competed for a mate. Our findings suggest that the microbiota is a key determinant of male mosquito life history with direct implications for optimizing production of males for control programs.

**Teaser:** The microbiota of *Aedes aegypti* mosquitoes affects multiple male life history traits.

## Introduction

The yellow fever mosquito, *Aedes aegypti*, transmits several human pathogens including viruses such as dengue, Zika, chikungunya, and yellow fever, which collectively cause hundreds of millions of cases and tens of thousands of deaths each year in tropical and subtropical regions (CDC, 2016; Powell, 2018; Brady & Hay, 2020; WHO, 2023; WHO, 2024). Effective control of *Ae. aegypti* populations is essential to reduce the burden of these diseases (Benelli et al., 2016; McGregor & Connelly, 2021; Lorenz & Chiaravalloti, 2022). Traditional methods, including environmental management, larval source reduction, and use of insecticides play key roles in managing mosquito populations (Benelli et al., 2016; McGregor & Connelly, 2021; Lorenz & Chiaravalloti, 2022). However, increasing resistance to insecticides underscores the need for novel strategies (Liu, 2015; CDC, 2024).

Promising alternative approaches for mosquito control include Sterile Insect Technique (SIT) and Incompatible Insect Technique (IIT), and global utilization of these approaches has increased dramatically in recent years (Harris et al., 2012; Kittayapong et al., 2019; Crawford et al., 2020; Aldridge et al., 2024; Martín-Park et al., 2024; Morreale et al., 2025). SIT relies on sterilizing males, e.g. through genetic methods or irradiation, before releasing them into the wild, where they mate with wild females, leading to non-viable offspring (Alphey et al., 2010; Lees et al., 2015; Dyck et al., 2021; Aldridge et al., 2024). This method has been successfully implemented in multiple vector control programs, including for *Ae. aegypti* (Harris et al., 2012; Aldridge et al., 2024; Morreale et al., 2025). IIT utilizes the endosymbiotic bacterium *Wolbachia*, which induces cytoplasmic incompatibility, preventing wild females from producing viable offspring when they mate with *Wolbachia*-infected males (Landmann et al., 2009). IIT has also been successfully used in recent years to control *Ae. aegypti* populations, either alone or in combination with the use of sterilizing radiation (Kittayapong et al., 2019; Crawford et al., 2020; Martín-Park et al., 2024). Both techniques require the mass rearing and release of millions of modified male mosquitoes that can successfully mate with wild females. Thus, a better understanding of factors impacting the physiology of male mosquitoes could serve to improve the efficacy and efficiency of these programs (Helinski & Harrington, 2013).

The mating performance of male *Ae. aegypti* mosquitoes plays a crucial role in population suppression strategies (Helinski & Harrington, 2013; Qureshi et al., 2019; Cator et al., 2021; Wyer et al., 2024). Mating lasts only a few seconds in *Ae. aegypti*, and during this brief time, males must efficiently transfer sperm and seminal fluid to the female. Seminal fluid contains a cocktail of proteins, hormones, and other molecules that induce dramatic effects on female physiology, including making her refractory to re-mating for an extended period of time, often the rest of her life (Avila et al., 2011; Helinski et al., 2012). For SIT and IIT to be successful, released males must efficiently copulate and transfer ejaculate to wild females. Additionally, male longevity and starvation resistance are also key considerations for SIT male success (Benedict et al., 2009; Helinski & Harrington, 2013). Males are produced for SIT at centralized facilities and transported to release sites in containers with variable access to food (Guo et al., 2022). Males with higher starvation resistance may be more likely to survive transport or live longer in release sites with limited food availability. Similarly, increased male longevity will potentially require decreased numbers of releases, thus decreasing cost and difficulty of successful population control. Therefore, considering aspects of the mass rearing process that have the potential to impact the mating performance, longevity, and field competitiveness of released males represents a key strategy for effective suppression of *Ae. aegypti* populations and reducing the transmission of mosquito-borne diseases.

One largely unexplored aspect of male biology that has high potential relevance for adult male life history traits is the environmentally acquired microbiota. Numerous previous studies have shown that the mosquito microbiota plays a vital role in shaping the biology of *Ae. aegypti* mosquitoes, influencing fitness-related traits such as pupation success, development time, body size, longevity, blood meal digestion, and reproductive success (Coon et al., 2014; Dickson et al., 2017; Correa et al., 2018; Martinson & Strand, 2021; Giraud et al., 2022; LaReau et al., 2023; Harrison et al., 2023; Roman et al., 2024; Díaz et al., 2025). The absence of a microbiota during larval development results in a failure to develop (Coon et al., 2014) unless adequate dietary supplementation is provided (Correa et al., 2018; Wang et al., 2021). Bacteria acquired during the larval stage provide essential nutrients, including B vitamins, which are critical for metabolism but must be obtained from the diet or microbial symbionts (Coon et al., 2014; Martinson & Strand, 2021; Romoli et al., 2021; Wang et al., 2021; Romoli et al., 2024). Critically, nearly all research on the effects of the microbiota on adult mosquito life history traits has focused on adult female mosquitoes because only females transmit pathogens. Studies that include adult male *Ae. aegypti* are less common but have shown that the microbiota can impact adult male longevity (Harrison et al., 2023) and wing length (Correa et al., 2018; Martinson and Strand, 2021; Roman et al., 2024; Díaz et al., 2025). The microbiota may impact other life history traits in male *Ae. aegypti*, and further investigation into the effects of the microbiota on male mosquitoes is warranted.

In the current work, we present a comprehensive investigation into the impacts of the microbiota on multiple life history traits in male *Ae. aegypti* mosquitoes. We find that the microbiota impacts adult male longevity, starvation resistance, wing length, and copulation success in non-competitive mating scenarios. Moreover, we find that the microbiota is most impactful during larval development, as controlled removal of the microbiota at only the adult stage had no effect on adult longevity. Together, these findings provide novel and foundational insight into the role of the microbiota on the biology of male *Ae. aegypti*.

## Materials and methods

### Experimental design overview

We designed three experiments (Experiment 1, 2, and 3, described in detail below) to investigate the impact of microbiota modulation or microbiota absence on life history traits of unsexed larvae and adult male *Ae. aegypti*. In all experiments, we used Thailand (Thai) strain *Ae. aegypti* mosquitoes (Villarreal et al., 2018). After hatching larvae from sterilized eggs as described below, the axenic larvae were randomly allocated to one of three treatments according to the objectives of each experiment: axenic (AX), i.e. maintained in a microbiota-free state throughout the experiment; monoxenic (MX), i.e. inoculated with only *E. coli*; or lab community (LC), i.e. mosquitoes inoculated with an undefined microbiota derived from laboratory reared mosquitoes (Fig. 1).

**Fig. 1.**
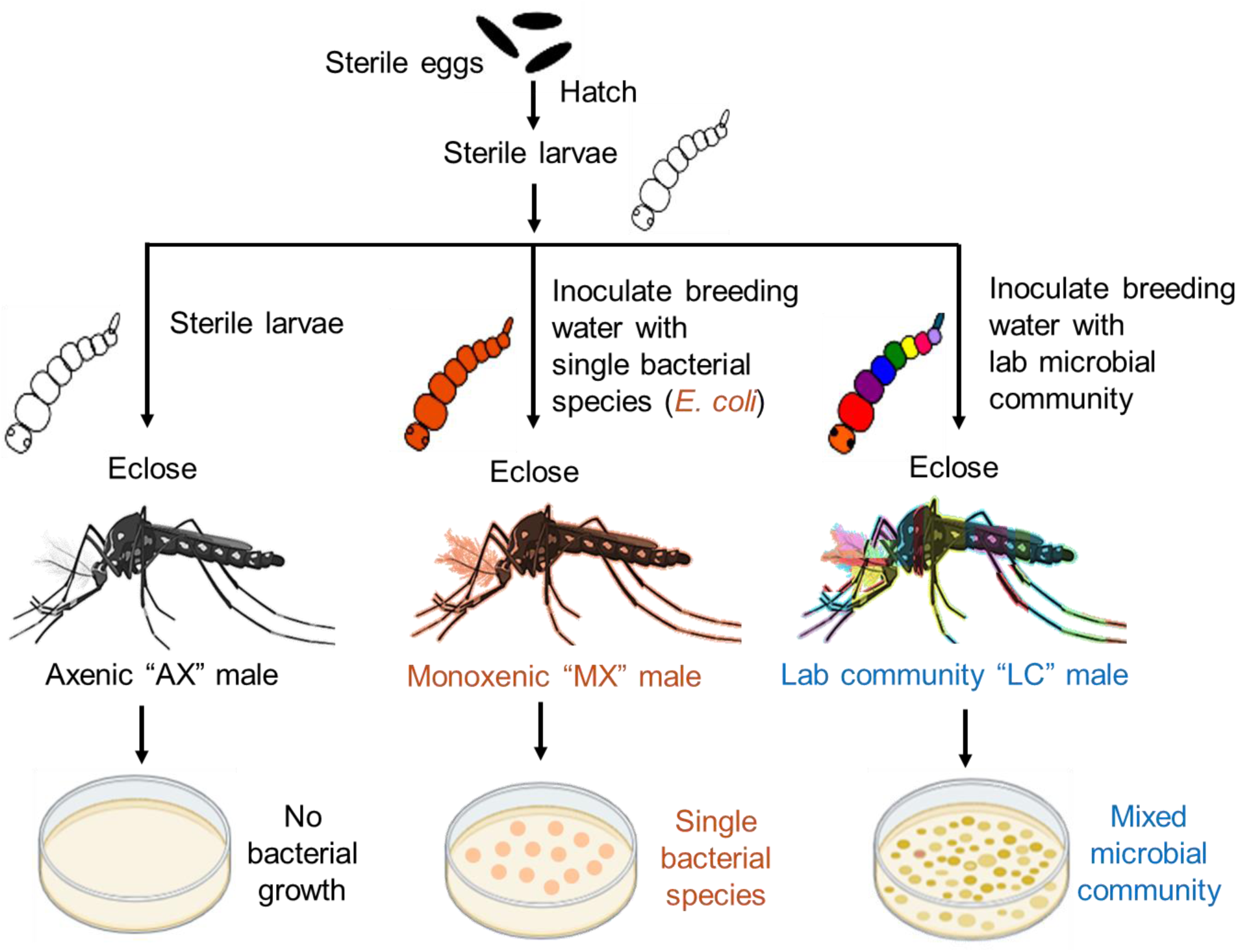
Experimental design overview. Three larval treatments were used throughout the study. Thai *Aedes aegypti* eggs were sterilized and hatched under sterile conditions and the resulting axenic larvae were randomly allocated to one of three treatment groups. The first group remained bacteria-free (axenic, AX), the second group was inoculated with only *E. coli* (monoxenic, MX), and the third group was exposed to a laboratory microbial community (lab community, LC). The success of our sterilization procedure and sterile rearing conditions was initially validated by PCR targeting the bacterial 16S rRNA gene and culturing larval water and adult males on LB agar (Fig. S1, Fig. S2). Success of each microbiota treatment was validated during each experiment by culturing homogenized adult males on LB agar plates (Fig. S3). This image was created using BioRender software (https://www.biorender.com/).

### Mosquito egg sterilization

Mosquito eggs were sterilized as in Coon et al., (2014) with some modification. A piece of filter paper containing *Ae. aegypti* Thai eggs, aged between one week to three months, was sterilized as follows: in a petri dish, the filter paper was rinsed with 70% ethanol, and the eggs were gently scraped using a sterile metal spatula and transferred to a sterile 15 mL falcon tube. The eggs were soaked in 70% ethanol for 5-7 minutes, then the ethanol was discarded. Next, a solution of 3% bleach with 0.1% of 25H Quat disinfectant cleaner (3M, USA) was added to the tube for 5-7 minutes, followed by removal of the bleach solution. The eggs were then immersed again in 70% ethanol for 5-7 minutes, then the ethanol was discarded. To remove any residual bleach or ethanol, the eggs were then rinsed three times with sterile deionized (DI) water. The sterile eggs were placed in a sterile petri dish with sterile 1X PBS and left to hatch in the biosafety cabinet. All procedures took place inside a biosafety cabinet using autoclaved tools.

### Preparation of axenic larval diet

Larvae were reared with sterile heat-killed *Escherchia coli* K12 agar plugs (HK-*E. coli* food plugs) as described in Correa et al., (2018) and Hyde et al., (2019). The HK-*E. coli* plugs were prepared as follows: the day before food preparation, two 500 mL flasks of sterile LB broth were inoculated with *E. coli* K12 and incubated overnight at 37°C with shaking. After 24 hours of incubation, the *E. coli* was harvested by centrifugation at 10,000 rpm for 10 minutes, and the resulting pellets were resuspended in 20 mL of sterile 1X PBS. For the food preparation, 3 g of liver powder (Now Foods, USA) were mixed with 2 g of brewer’s yeast powder (Solgar, inc., USA) in experiments 1 and 3 or 2 g of granulated yeast extract (Fisher Scientific, USA) in experiment 2. Then 2 g of the Liver:Yeast (LY) mixture, combined with 1 g of agar (Fisher Scientific, USA), were added to 40 mL of DI water to form an LY/agar mixture. The 20 mL bacterial suspension was then added to the 40 mL LY/agar mixture, and the entire mixture was autoclaved for 1 hour. Once autoclaved, the mixture was poured into three sterile petri dishes (20 mL / petri dish) inside a biosafety cabinet and allowed to solidify. We used sterile 1.2 mL cryovials (Fisher Scientific, USA) to excise cylindrical plugs from the solidified agar mixture. The food plugs were stored in the dark at 4°C and used for a maximum of three weeks after preparation.

### Preparation of bacteria for introduction to axenic mosquitoes

Two strains of *E. coli* were used for introduction to axenic mosquitoes (*E. coli* K12 MG1655, and *E. coli* S17 pPROBE-mCherry). Use of each strain is detailed for each experiment below. *E. coli* S17 pPROBE-mCherry (*E. coli* S17) contains a kanamycin resistance cassette. To prepare inocula for the MX treatment, *E. coli* was grown from a single colony in liquid LB (plus 50µg/mL kanamycin for *E. coli* S17) at 30°C with shaking for 18-20 hours. Cultures were washed two times with sterile 1X PBS and resuspended in sterile 1X PBS to OD_600_ = 1.0. To prepare inocula for the LC treatment, we homogenized one fourth-instar larva, conventionally reared (CN, i.e. reared in our laboratory colony with no manipulation of the microbiota), in 500 µL of DI water. A fresh homogenate was prepared for each replicate of each experiment and used to inoculate all wells/flasks for that experiment.

### Maintaining larvae, pupae and adults

Larvae were provided with sterile food (see “Preparation of axenic larval diet” above for details). Rearing conditions were maintained at 27°C ± 1, with 70-80% relative humidity. In order to permit axenic larval development, larvae were reared without light (Hyde et al., 2019), and this was applied consistently across all treatments and experiments. As pupae emerged and adults eclosed, individuals were transferred to containers under sterile conditions inside a biosafety cabinet or laminar flow hood. As pupae emerged, they were collected using sterile transfer pipets, individually placed in sterile 1.5 mL tubes with small holes in the bottom of the tube for ventilation, and placed cap-side down until eclosion. Upon emergence, adult male mosquitoes were transferred to sterile 8oz or 16oz Mason jars (Ball) fitted with mesh covers and provided with a filter-sterilized 10% sucrose solution. Then the jars were inserted into sterile 86 oz polypropylene plastic tubs (Aaron Packaging, Inc.), which were sealed with sterile lids to maintain sterile conditions. Pupae and adults were maintained under sterile conditions at 27°C ± 1, with 70-80% relative humidity and a 14-hour light/10-hour dark cycle.

### Verification of sterility of axenic larvae and adults

To initially verify that our axenic protocol successfully eliminated bacteria and maintained sterile conditions, we performed PCR amplification of the bacterial 16S rRNA gene using primers fD1/fD2 and rP1/rP2 (Weisburg et al., 1991) with DNA extracted from axenic larvae and axenic adult males as template using the DNeasy Blood & Tissue kit (Qiagen, USA) according to the manufacturer’s instructions. PCR reagents were prepared as follows: 0.5 μL of each primer at a concentration of 10 μM was added to 12.5 μL of MyTaq Red master mix (Fisher Scientific, USA), and then molecular grade water was added to a final volume of 25 μL. The PCR cycling conditions were as follows: 95°C for 3 minutes, followed by 30 cycles of (95°C for 30 seconds, 55°C for 30 seconds, 72°C for 30 seconds), then 72°C for 10 minutes, and finally a 10°C hold. Following gel electrophoresis, an absence of visible bands in axenic individuals confirmed the absence of microbial contamination (Fig. S1). We also initially verified our axenic protocol by culturing larval water and adult males on LB agar (Fig. S2). Additionally, for every experiment, three adult males from each treatment were individually homogenized in 150 µL of sterile 1X PBS. Then, 100 µL of the homogenate was cultured on LB plates and incubated at room temperature for one week to confirm the microbiota treatment (Fig. S3).

### Experiment 1: Comparing development time and adult male life history traits between MX and LC individuals

In this set of experiments, we examined how adult males colonized by a single bacterial isolate (MX) differ in life history traits compared to those reared with a laboratory microbial community (LC). For each replicate, approximately 150 axenic larvae were distributed across six sterile 25 cm^2^ cell culture flasks, each containing 25 mL of sterile DI water (25 larvae per flask). Each flask was supplied with three HK-*E. coli* food plugs, prepared using liver powder and brewer’s yeast powder (see “Preparation of axenic larval diet” above). Three flasks were inoculated with 1 µL of *E. coli* S17 at OD_600_ = 1.0, resulting in an inoculating dose of ∼5 × 10^4^ CFU/mL, forming the MX group. The remaining three flasks were inoculated with 5 µL of a laboratory microbial community preparation (see “Preparation of bacteria for introduction to axenic mosquitoes” above), resulting in an inoculating dose of ∼5 × 10^3^ CFU/mL, forming the LC group. Inoculating dose was determined for both treatments by either serially diluting and culturing larval water on LB agar immediately following inoculation or culturing an inoculum dilution series on LB agar. We used a lower inoculating dose for LC because we observed that concentrations above 5 × 10^3^ CFU/mL had a lethal effect on the first-instar larvae using this rearing protocol. We note that the CFU/mL estimate for LC may be an underestimate because not all members of the laboratory community are expected to grow equally well on LB.

Pupation and eclosion rates were recorded for all individuals from each treatment. Adult male mosquitoes were collected within two days of eclosion, placed in a sterile container, and provided either 10% sterile sucrose to assess longevity or water only to assess starvation resistance. Individual male survival was recorded daily until all individuals in each group had died. The experiment was conducted in four replicates for pupation and eclosion, totaling 49-75 larvae per treatment per replicate (total *n* = 249-275 per treatment) and three replicates for survival, totaling 14-17 adult males per treatment per replicate (total *n* = 46 per treatment in sugar-fed survival and *n* = 45 for water-fed survival).

In addition, we monitored the effect of treatment (MX or LC) and the presence of larvae on bacterial CFU in rearing water at multiple timepoints. To do this, we sampled 100 µL of water from MX and LC flasks with and without larvae at 2, 4, 6, and 8 days after introducing *E. coli* S17 or the laboratory microbial community. Samples were serially diluted tenfold up to 10^−6^, and 100 µL from each dilution was spread on LB agar and incubated at room temperature for one week to allow growth. CFUs were then counted for each sample and back-calculated to determine CFU per mL. Bacterial load data were collected from three independent replicates.

Furthermore, to evaluate the effect of the microbiota on male mating-related traits, we conducted experiments under non-competitive and competitive mating scenarios (Lang et al., 2018) (Fig. 2). For each scenario, 4-5-day-old MX and LC virgin males were used along with age-matched, conventionally reared (CN) virgin females.

**Fig. 2.**
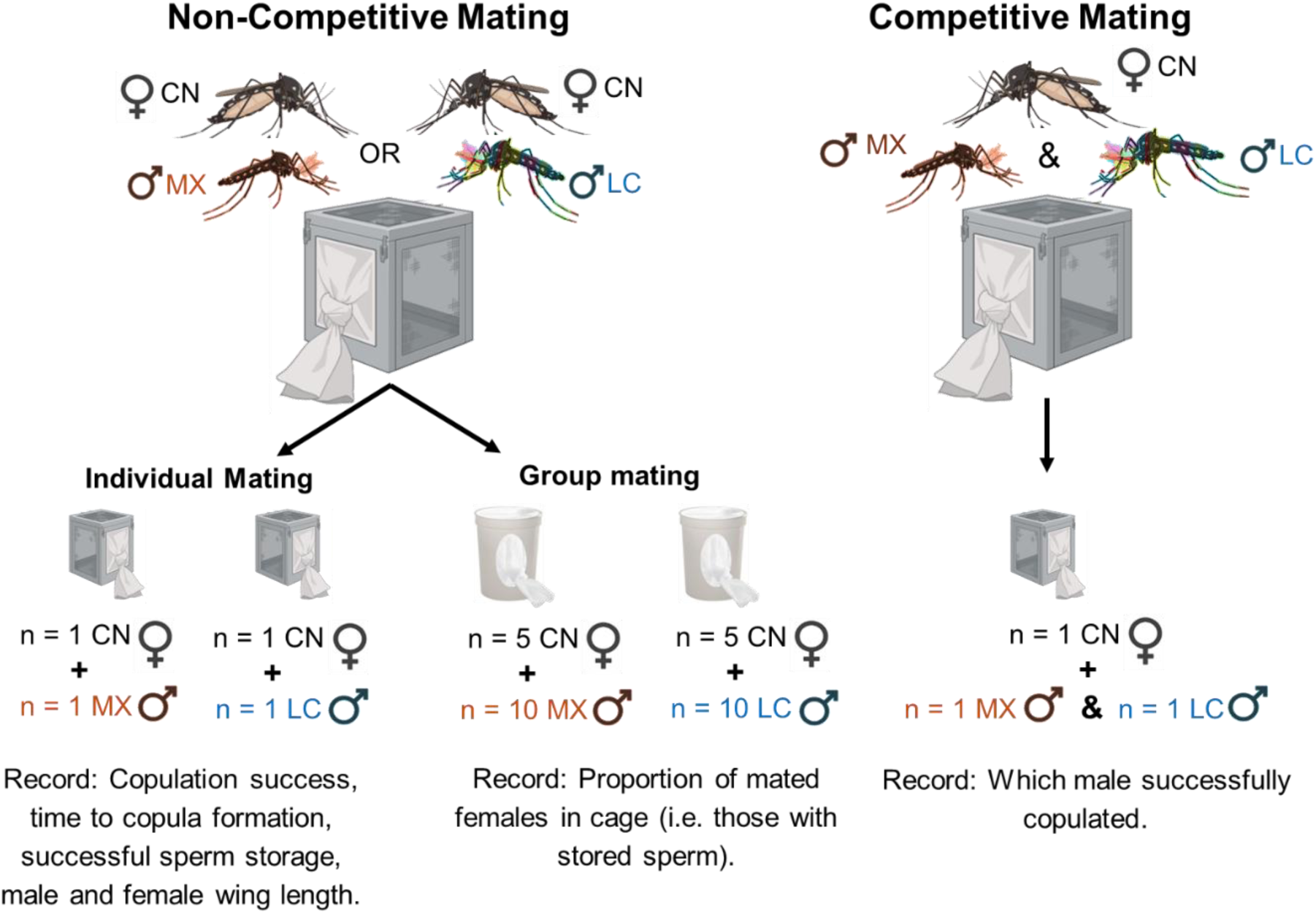
Overview of experimental design for mating experiments in Experiment 1. MX and LC male mating-related traits were assessed in two different scenarios: non-competitive and competitive mating. In the non-competitive scenario, males from either the MX or LC group were allowed to mate with virgin conventional (CN) females either individually, i.e. one male with one female, for up to 5 minutes, or in groups, i.e. ten males with five females, for 30 and 90 minutes. The former is termed “Non-competitive individual mating” and the latter “Non-competitive group mating.” In the competitive mating scenario, one MX male and one LC male were aspirated into a cage simultaneously and allowed to compete for a single virgin CN female for a maximum of 15 minutes. This is termed “Competitive mating”. This image was created using BioRender software (https://www.biorender.com/).

For non-competitive individual mating, an individual trial involved placing one MX or one LC male in a metal cage (8 x 8 x 8 in; 32 mesh Lumite®/vinyl window, BioQuip, USA) with a single, age-matched virgin CN female. Each pair was allowed to attempt to mate for a maximum of 5 minutes, with the cage gently shaken every 30 seconds to encourage flight. We recorded whether copulation took place within the 5 minutes window as well as the time in seconds from when the trial started until copula formation occurred. Afterward, each female’s spermathecae were dissected under a dissecting microscope and examined under a compound microscope to determine whether sperm storage was successful. This experiment was conducted in four replicates, with each replicate involving ten trials. After completion of the mating experiment, images of MX and LC male and female mosquito wings were taken using a dissecting microscope. The wing length was measured using Fiji ImageJ software, by drawing a straight line from the alular notch to the wing apex, representing the maximum wing length (Bock and Milby, 1982). Length measurements were determined after setting a scale using a micrometer. Males used in this experiment arose from the same cohort as the individuals used in the pupation, eclosion, and adult male survival measurements described above. Their pupation and eclosion data were included in the pupation and eclosion analyses described above, but they were discarded after mating trials and were therefore excluded from longevity measurements.

For non-competitive group mating, an individual trial involved placing ten LC or MX males in a 86 oz polypropylene plastic tub (Aaron Packaging, Inc.) fitted with cotton stockinette netting (Medline, USA), with five age-matched virgin CN females and allowing them to mate for either 30 or 90 minutes. At the end of each trial, the females from each group were removed, and each female’s spermathecae were dissected and examined under a compound microscope to determine whether sperm storage was successful. The number of mated females (determined by the presence of sperm in the spermathecae) was recorded for each trial, and the experiment was conducted in six replicate trials for each treatment/time point combination.

For competitive mating, an individual trial involved placing one MX male and one LC male together in a metal cage, then introducing one virgin CN female to initiate mating competition between the two males. Timing began upon the female’s entry, and the cage was shaken every 30 seconds to encourage flight. Each trial lasted a maximum of 15 minutes. When one male formed a copula with the virgin CN female (the “winning” male), the time was recorded, and the mating pair was removed using an aspirator and immediately transferred to a clean falcon tube, while the non-mated male (the “losing” male) was placed in a separate tube. Both mated and non-mated males were surface sterilized with 70% ethanol for 30 seconds, washed twice with sterile water, homogenized in 150 µL of sterile 1X PBS, and 100 µL of the homogenate was cultured on LB agar to determine the identity of the winning male. In every trial, one homogenate yielded very few colonies, and the other homogenate yielded relatively high numbers of colonies composed of multiple different colony types. We ascribed each of the former to the MX treatment and each of the latter to the LC treatment (Fig. S4). This experiment was conducted in three replicates, with each replicate comprising 10-12 trials.

### Experiment 2: Comparing development time, adult male longevity, and male wing length among AX, MX and LC individuals

In this experiment, we investigated the impact of the complete absence of a microbiota on larval development time, adult male wing length, and adult male longevity. Three experimental treatments were applied to larvae: AX, MX, and LC. After egg sterilization, axenic larvae were transferred to nine sterile 6-well plates (2-3 larvae per well) with each well containing 5 mL of sterile DI water and one HK-*E. coli* food plug prepared using liver powder and granulated yeast extract (see “Preparation of axenic larval diet” above). Three of the 6-well plates remained AX. Three were inoculated with 1 µL of *E. coli* S17 at OD_600_ = 1.0, resulting in an inoculating dose of ∼2 × 10^5^ CFU/mL (determined by culturing an inoculum dilution series on LB agar), forming the MX group. The other three were inoculated with 5 µL of a laboratory microbial community preparation (see “Preparation of bacteria for introduction to axenic mosquitoes” above), resulting in an inoculating dose of ∼2 × 10^4^ CFU/mL (determined by culturing an inoculum dilution series on LB agar), forming the LC group. Pupation and eclosion rates were recorded for all individuals from each treatment. Adult male mosquitoes were collected within two days of eclosion, transferred to sterile containers, and provided with sterile cotton soaked in a sterile 10% sucrose solution. Survival was monitored daily until all males died. Wings were collected from males at the time of death for wing length measurement as described in Experiment 1. This experiment was conducted in three replicates for pupation and eclosion; there were three 6-well plates per replicate, totaling 24-36 larvae per treatment per replicate (total *n* = 72-108 per treatment). Survival and wing length were measured in two subsequent replicates; 10-15 adult males were measured per treatment per replicate (total *n* = 22-25 per treatment). Total sample size for wing length measurement was 5-12 per treatment per replicate (total *n* =17-19 per treatment).

### Experiment 3: Assessing the effect of the adult microbiota on adult male longevity

In this experiment, we used ampicillin treatment and *E. coli* feeding to assess how differences in the microbiota specifically at the adult stage impact adult male longevity. To do this, we used MX larvae to create four adult male treatment groups: 1) AX_Amp_ was generated by treating MX fourth instar larvae with ampicillin shortly prior to pupation to eliminate the microbiota; 2) AX_Amp+*E. coli*-fed_ was generated identically to AX_Amp_, then exposed to *E. coli* by feeding as adults; 3) MX is identical to previous experiments; and 4) MX*_E. coli_*_-fed_ were MX that were additionally exposed to *E. coli* by feeding as adults.

To generate these four treatment groups, approximately 150 axenic larvae were transferred into ten sterile 25 cm^2^ cell culture flasks (15 larvae per flask), each containing 25 mL of sterile DI water. All flasks received three HK-*E. coli* food plugs prepared using liver powder and brewer’s yeast powder (see “Preparation of axenic larval diet” above) and were inoculated with 1 µL of *E. coli* K12 MG1655 at OD_600_ = 1.0, resulting in an inoculating dose of ∼ 5 × 10^4^ CFU/mL (determined by culturing an inoculum dilution series on LB agar). *E. coli* K12 MG1655 was used for this experiment because it is known to be highly susceptible to ampicillin (DSMZ Institute). At the late 4^th^ instar stage, one day before pupation, five flasks were treated with 200 µg/mL of ampicillin to eliminate *E. coli* and generate AX males, termed here AX_Amp_, and five other flasks remained untreated to generate MX males. Pupation rates were recorded for both treated and untreated groups, and pupae from both groups were collected 2-3 days post-pupation. Pupae from the ampicillin-treated group were rinsed twice with sterile DI water and exposed again to 200 µg/mL of ampicillin. The pupae with the ampicillin treated DI water were then transferred to 1.5 mL tubes until eclosion. Males were collected within two days of eclosion, split into four sterile glass jars (two jars per treatment, 10-15 males per jar), and maintained with sterile cotton plugs soaked in 10% sterile sucrose solution. The success of ampicillin treatment was validated by culturing homogenized ampicillin treated and untreated adult males on LB agar plates (Fig. S5).

The day before *E. coli* feeding, all males were starved and deprived of water by removing the cotton plugs. After 24 hours of starvation, one jar of AX_Amp_ and one jar of MX were provided with *E. coli* K12 MG1655 prepared by diluting a OD_600_ = 1.0 culture ten-fold in sterile 10% sucrose, resulting in a final concentration of ∼1 × 10^8^ CFU/mL. This generated the AX_Amp+*E. coli*-fed_ and MX*_E. coli_*_-fed_ treatment groups. The two other jars were provided with sterile cotton plugs soaked in sterile sucrose solution, and this generated the AX_Amp_ and MX treatment groups. After 24 hours of feeding, the *E. coli*-treated cotton plugs were replaced with sterile cotton plugs soaked in sterile 10% sucrose. Male survival was monitored and recorded until all individuals in each group had died. This experiment was replicated thrice; pupation and eclosion data were recorded for all three replicates, and survival was recorded for two replicates. Sample sizes were 45-75 larvae per treatment per replicate (total *n* = 195-225 per treatment) and 9-13 adult males per treatment per replicate (total *n* = 20-25 per treatment).

### Statistical analysis

Analyses were performed in R (R Core Team, 2025) and R studio (Posit team, 2025) using packages survival (Therneau and Grambsch, 2000; Therneau, 2023), survminer (Kassambara et al., 2021), car (Fox & Weisberg, 2019), ggplot2 (Wickham, 2016), interactions (Long, 2024), arm (Gelman & Su, 2024), stats (R Core Team, 2025), dplyr (Wickham, et al., 2023), tidyverse (Wickham et al., 2019), ggpubr (Kassambara, 2023a), rstatix (Kassambara, 2023b), lavaan (Rosseel, 2012), and DHARMa (Hartig, 2025).

For survival analyses, we fit Cox Proportional Hazards models using treatment and replicate as predictors. Following model fitting, we performed an Analysis of Deviance to determine which variables in the model significantly predicted survival. In instances where there were more than three levels for treatment, we used the pairwise_survdiff() function from the survminer package to perform pairwise comparisons between treatment groups. For survival data in Experiment 3 and most pupation and eclosion datasets, the data did not conform to the assumptions of proportional hazards. In these instances, we used a logrank test in the survdiff() function from the survival package to determine the overall effect of treatment and the pairwise_survdiff() function from the survminer package to perform pairwise comparisons.

To analyze wing length data, we fit linear models using treatment and replicate as predictors. We then performed an ANOVA to assess the overall significance of each predictor variable and used Tukey’s HSD test to evaluate pairwise comparisons within treatment. To analyze bacterial load data, we fit a mixed measures ANOVA testing the effect of day as a repeated measure within flask and effects of bacterial treatment (MX or LC) and larvae (present or absent) as independent predictor variables that were not repeated measures. Interaction plots were generated using the interaction.plot() function from the *stats* package and the interact_plot() function from the *interactions* package.

For non-competitive individual mating, we evaluated time to copula formation in seconds (continuous response variable) and copulation success (binary response variable). To analyze time to copula formation data, we fit a general linear model (GLM) using a Gaussian distribution with treatment, female wing length, male wing length, and replicate as predictor variables. A Type III ANOVA was then used to assess the significance of each model term. The full model assessed a 3-way interaction between treatment, female wing length, and male wing length, all potential 2-way interactions, and main effects of each predictor. To analyze copulation success data, we fit a GLM using a binomial distribution with treatment, female wing length, male wing length, and replicate as predictor variables. A Type III ANOVA was then used to assess the significance of each model term. The full model assessed a 3-way interaction between treatment, female wing length, and male wing length, all potential 2-way interactions, and main effects of each predictor. For each of these variables (time to copula formation and copulation success), we also performed a path analysis to examine causal relationships among the microbiota treatment and male mosquito wing length outlined in the conceptual models (Fig. S6A, Fig. S7A) to identify indirect pathways mediating time to copula formation and copulation success. Predictor variables included in the model tested the direct effects of microbiota treatment and male wing length on time to copula formation and copulation success, and the indirect effects of microbiota treatment via male wing length. Because time to copula formation was recorded only when copulation was successful, separate path analysis models were run for time to copula formation and copulation success due to unequal sample sizes. Variables were assessed for assumptions of normality and checked to ensure no multicollinearity (r^2^ < 0.7) (Grewal et al., 2004). Time to copula formation was scaled to address high variance and model convergence issues. Maximum likelihood methods were used for parameter estimation (Fan et al., 2016). Copulation success was specified as categorical variable using the ordered argument and diagonally weighted least squares methods were used for parameter estimation (Li, 2021). Coefficient estimates of indirect effects were generated using 1000 bootstrap replicates. Model performance was evaluated using three standard metrics: 1) chi-square-based goodness-of-fit test with p > 0.05 indicating a model structure consistent with the data; 2) comparative fit index (CFI) with values of > 0.97 indicating acceptable model fit; and 3) root mean square error of approximation (RMSEA) with values < 0.05 indicating acceptable model fit (Schermelleh-Engel et al., 2003; Shipley, 2016).

To analyze non-competitive group mating data, we fit a GLM using a binomial distribution with treatment, time allowed to mate (30 or 90 minutes), and replicate as predictor variables. The full model included a 2-way interaction between treatment and time allowed to mate as well as the main effects of all predictors.

To analyze competitive mating data, we used a Chi-Square test to evaluate whether our observed data deviated from what would be expected under random chance (i.e. a 50/50 ratio between treatment groups).

Raw data, R code, and model outputs are available in File S1 and File S2.

## Results

### A monoxenic *E. coli* microbiota slows larval development and enhances adult male longevity and starvation resistance compared to a laboratory microbiota

We reared MX (monoxenic, i.e. inoculated with only *E. coli*) and LC (i.e. inoculated with an undefined microbiota derived from laboratory reared mosquitoes) individuals and assessed pupation, eclosion, adult male longevity, and adult male starvation resistance (i.e. survival when deprived of sucrose and provided only water) (Fig. 3A to 3D). Our results indicate that MX larvae had a slower pupation rate and delayed eclosion compared to the LC group when measured from the day of hatching (Fig. 3A and 3B; pupation rate, *P* < 2 × 10^−16^; eclosion rate, *P* < 2 × 10^−16^). Adult MX males had significantly extended longevity (Fig. 3C, *P* = 4.25 × 10^−7^) and significantly improved starvation resistance (Fig. 3D, *P* = 0.012) in comparison to LC males.

**Fig. 3.**
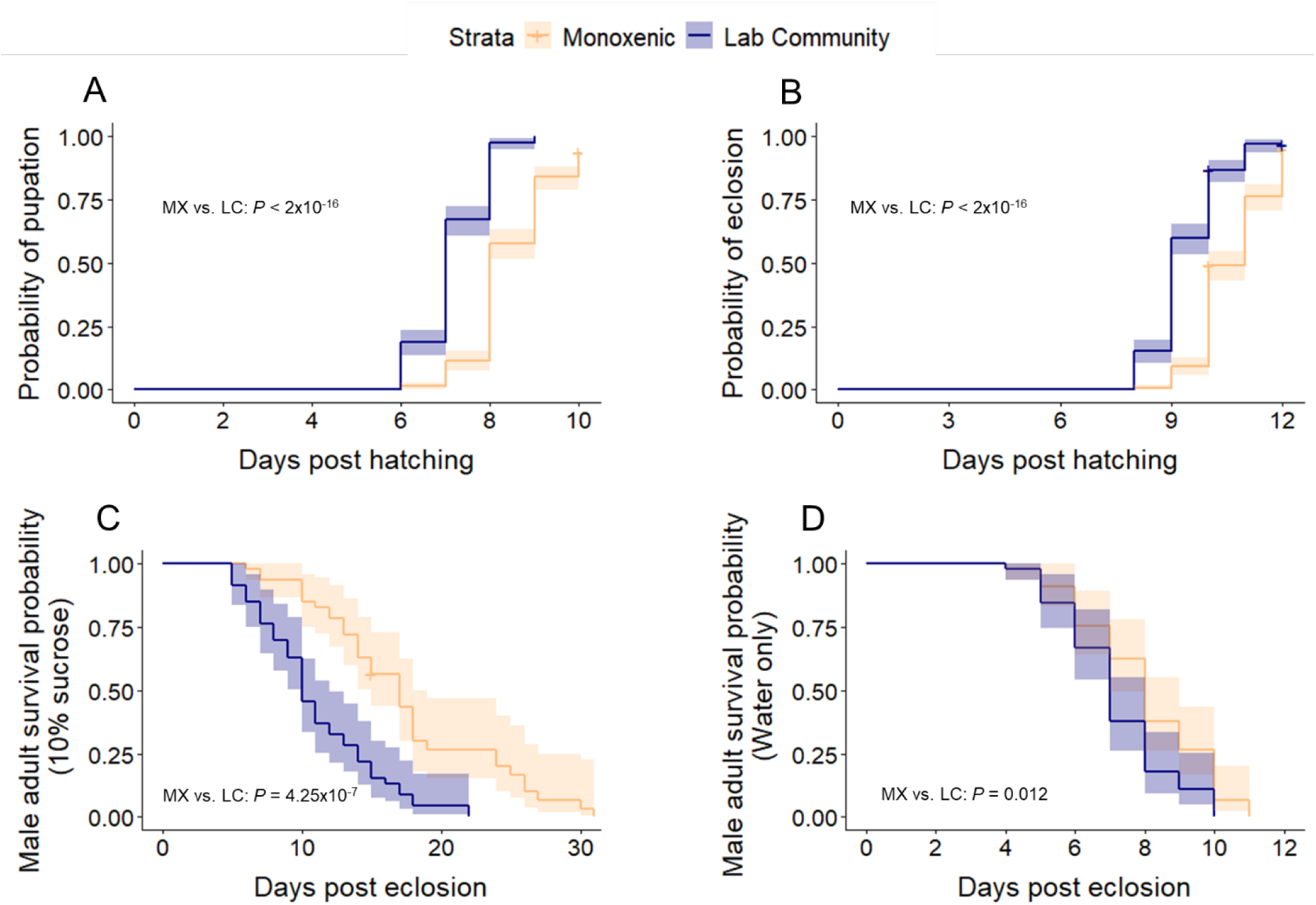
MX microbiota slows larval development and enhances adult male longevity and starvation resistance compared to an LC microbiota. Pupation and eclosion rates of larvae reared monoxenically with *E. coli* S17 (MX) were compared to those of larvae reared with the laboratory microbial community (LC). Following eclosion, adult male longevity was assessed under two conditions: a standard sugar diet (to evaluate longevity), and water only (to evaluate resistance to starvation). Curves were generated using the Kaplan Meier method, and shaded areas around curves represent 95% confidence intervals. MX larvae showed a slower (A) pupation rate (*P* < 2 × 10^−16^) and (B) eclosion rate (*P* < 2 × 10^−16^) when measured from the day of hatching. (C) Longevity of adult males was also significantly affected, as MX males survived longer than LC males when fed a normal sugar diet (*P* = 4.25 × 10^−7^). (D) MX males exhibited significantly higher survival when only fed water (i.e. starvation resistance) than LC males (*P* = 0.012). Data were collected from four replicate experiments for pupation and eclosion, totaling 49-75 larvae per treatment per replicate (total *n* = 249-275 per treatment) and three replicate experiments for survival, totaling 14-17 adult males per treatment per replicate (total *n* = 46 per treatment in sugar-fed survival and *n* = 45 for water-fed survival).

### A monoxenic *E. coli* microbiota improves the mating success of male mosquitoes in some mating scenarios compared to a laboratory microbiota

MX adult males and LC adult males were allowed to mate with aged-matched CN females in both non-competitive and competitive (i.e. MX versus LC) mating scenarios. For non-competitive mating, we tested both individual mating (i.e. one male with one female) and group mating (i.e. five females with ten males from a single microbiota treatment group).

In an individual mating, non-competitive scenario, one male from either treatment group was placed in a cage with one CN female for five minutes. Once copulation occurred, the time was recorded, the female was examined for successful sperm storage, and the wing length of both the male and female was recorded. Of 54 copulation events (observed over 80 trials), 50 resulted in successful sperm storage (92.3% success rate), indicating that copulation success was generally representative of successful insemination. We found that MX males had significantly smaller wings compared to LC males (Fig. 4A, *P* = 1.50 × 10^−4^).

**Fig. 4.**
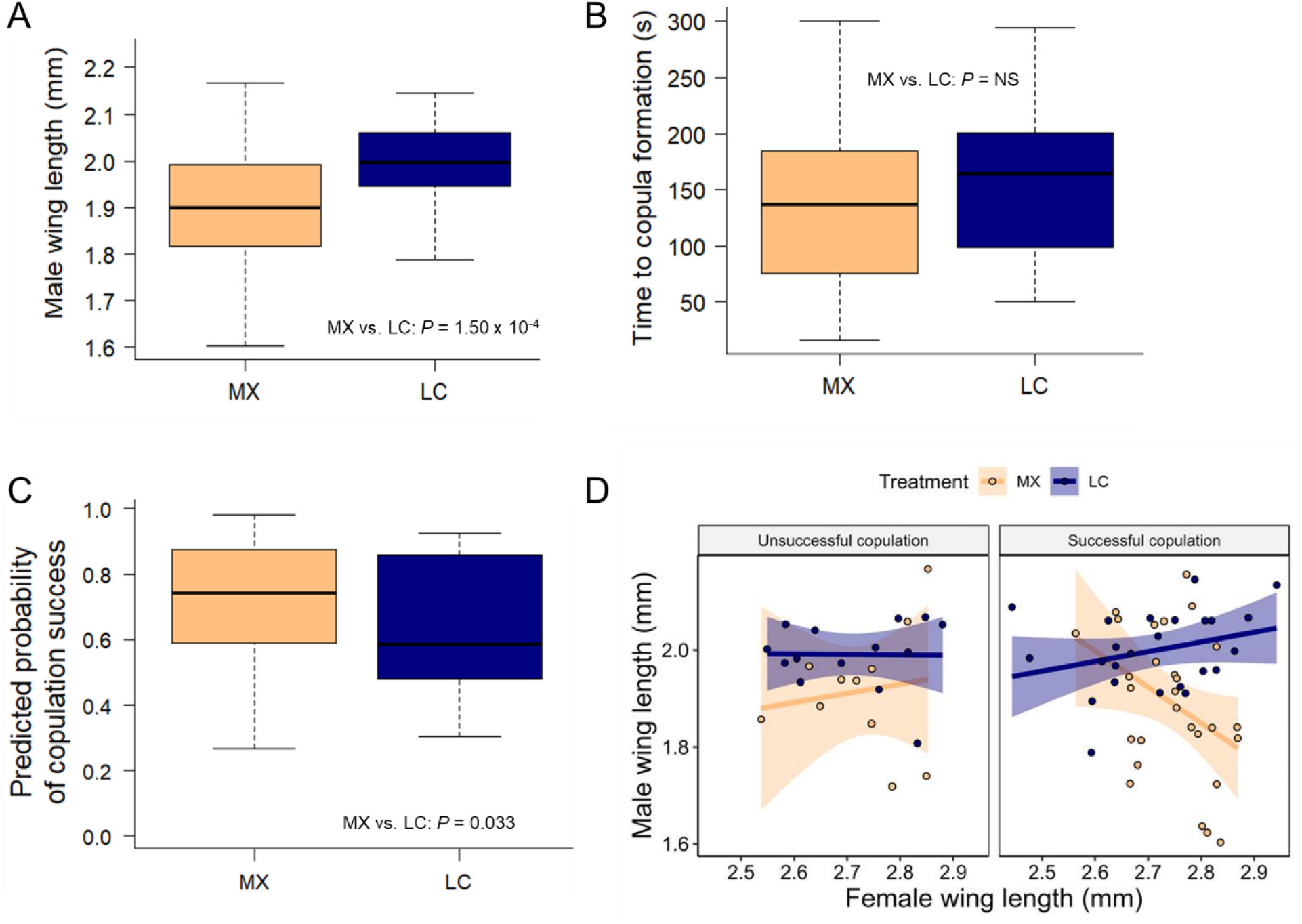
MX males have reduced wing length and increased likelihood of copulation success compared to LC males in an individual mating, non-competitive scenario. One MX male or one LC male was placed a cage with one CN female for five minutes. Copulation success was recorded, as was the time it took for copulation to occur, and the wing length of both the male and female used in each trial. Data were collected from four independent replicate experiments with *n* = 10 mating trials per treatment per replicate. (A) MX males had significantly smaller wings compared to LC males (*P* = 1.50 × 10^−4^). The box plot represents data collected from *n* = 40 males per treatment. (B) We observed no significant difference between MX and LC males for time to copula formation (*P* = 0.689). The box plot represents data collected from *n* = 53 trials. (C) MX males were significantly more likely to form a copula with a female compared to LC males (*P* = 0.033). The box plots display the variation in predicted probability of copulation success associated with each treatment; data were collected from n = 40 mating trials per treatment. (D) A linear regression analysis revealed a significant three-way interaction between treatment, female wing length, and male wing length (*P* = 0.034), indicating that the effect of the microbiota on copulation success differs depending on the body size of both the male and the female. Plotting mating trials parsed by copulation success illustrates that successful copulation (right panel) for MX is associated with a negative correlation between male and female wing length while LC is associated with a positive correlation between these traits. No such relationship was observed in either treatment for trials where copulation failed to occur (left panel).

We then performed analyses to understand the relationships between microbiota treatment, male wing length, and female wing length in the context of time to copula formation and copulation success. For time to copula formation, linear regression revealed no significant effects of any predictor (Fig. 4B). A path analysis showed no direct or indirect relationships between time to copula formation and microbiota treatment (Fig. S6B, Table S1).

For copulation success, linear regression revealed that MX treatment leads to a higher likelihood of copulation success compared to LC (*P* = 0.033, Fig. 4C), and path analysis revealed that the effect of microbiota treatment on copulation success is not mediated through the effect of microbiota on male wing length (Fig. S7B, Table S2). Furthermore, linear regression analysis revealed a significant three-way interaction between treatment, female wing length, and male wing length, indicating that the effect of the microbiota on copulation success differs depending on the body size of both the male and the female (*P* = 0.034); small MX males were more likely to copulate when females were large, but small LC males were more likely to copulate when females were small (and vice versa for large males) (Fig. S8). When copulation was successful, MX male wing length was negatively correlated with female wing length and the opposite was observed for LC males (Fig. 4D). For trials where copulation failed to occur, there was no relationship between wing length in either treatment (Fig. 4D).

In a group mating, non-competitive scenario, either MX or LC males were allowed to mate with conventionally reared (CN) females in groups of 10 males and five females for 30 or 90 minutes (Fig. 5A and 5B). After 30 minutes of mating, MX males displayed significantly higher mating success than LC males (Fig. 5A, *P* = 4.14 × 10^−4^). However, in treatment groups given 90 minutes to mate, we detected no differences in mating success between MX and LC males (Fig. 5B, *P* = 1.00).

**Fig. 5.**
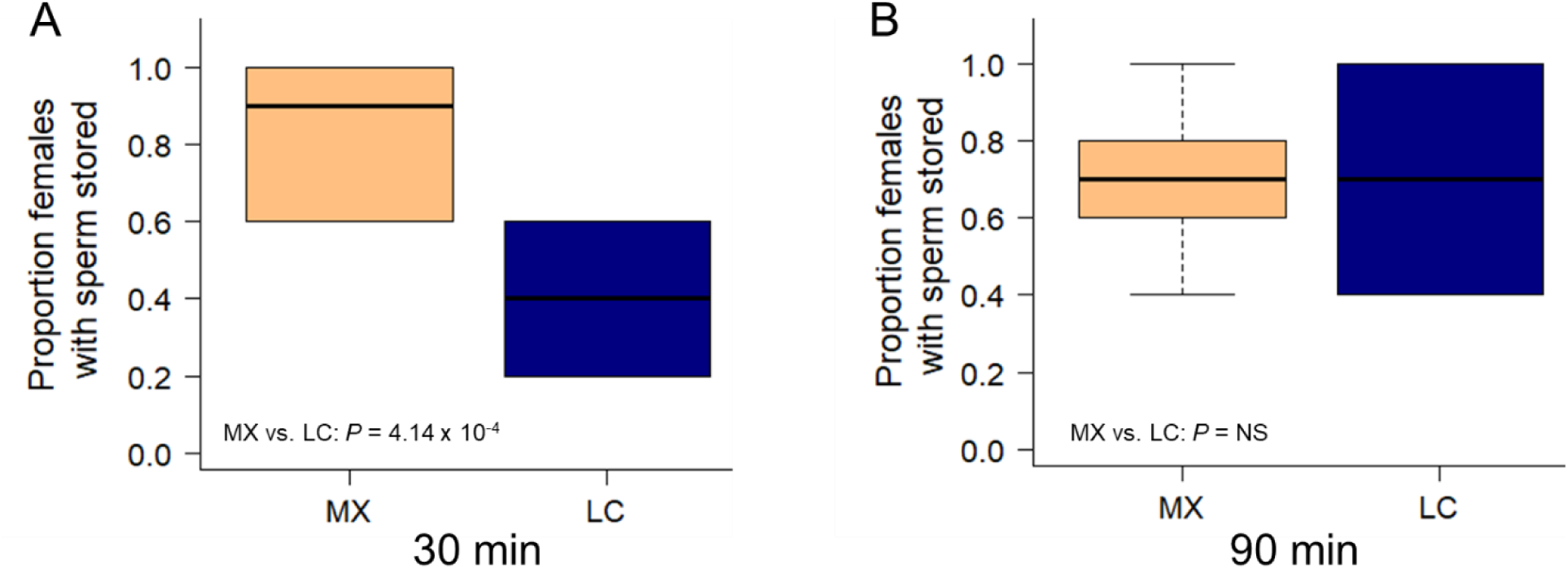
MX males have higher mating success than LC males in a group mating, non-competitive scenario. Either MX or LC males were allowed to mate with CN females in groups of ten males and five females for 30 or 90 minutes. The data were collected from six independent group mating trials. Box plots show the proportion of females from each trial (n = 6 proportion values per treatment) that had sperm in their spermathecae at the end of the mating period. (A) After 30 minutes of mating, females in cages containing MX males were significantly more likely to have mated than females in cages containing LC males (*P* = 4.14 × 10^−4^). (B) By 90 minutes, there was no significant difference in mating success between MX and LC (*P* = 1.00).

In a competitive mating scenario, one MX male and one LC male were placed together in a metal cage with a single virgin CN female and observed until one male successfully copulated with the female. We found that MX males had an equal ability to copulate as LC males when competing for a female (Χ^2^_1,32_ = 0.5, *P* = 0.478; Fig. 6A) despite some variability across replicates (Fig. 6B).

**Fig. 6.**
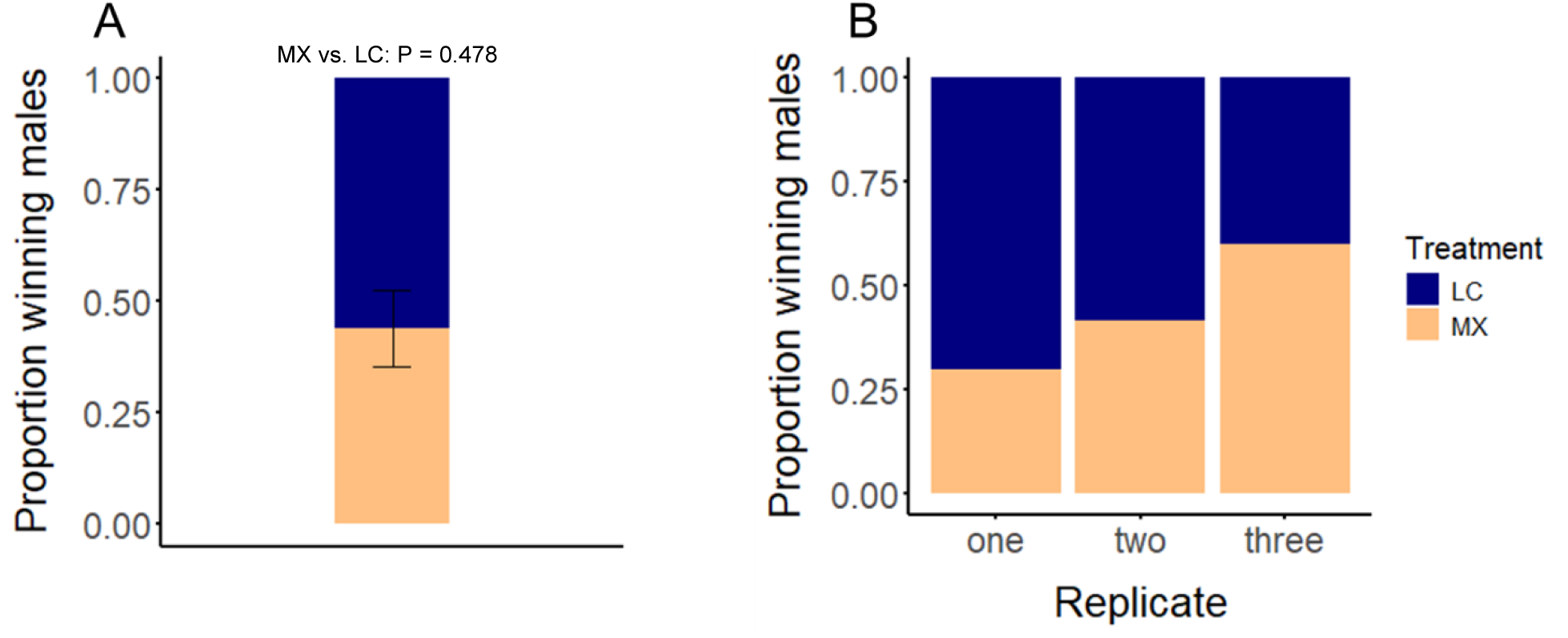
Microbiota treatment has no effect on copulation success in a competitive mating scenario. One MX male and one LC male were allowed to compete for one CN female, then the “winning” male (i.e. the male that successfully copulated with the female) was recorded. The experiment was replicated three full times, with each replicate consisting of 10-12 mating trials per replicate. (A) MX males are equally able to copulate when in competition with LC males. The figure shows the average proportion of trials from the three replicate experiments that were “won” by MX or LC treatment males (n = 3 proportion values). The error bar shows the standard error of the mean. A Chi Square test querying deviation of our observed results from an expected 50/50 ratio of success between treatments indicated no significant difference from the null hypothesis of equal success (Χ^2^_1,32_ = 0.5, *P* = 0.478). (B) Parsing the data by replicate shows that the effect of microbiota on competitive copula formation was variable across replicates.

### Bacterial load in larval rearing water differs between MX and LC and over time

In addition to examining the effects of microbial treatments on larval development and adult male traits, we investigated bacterial load in rearing water and whether larval presence influenced bacterial abundance. The initial concentration of *E. coli* S17 in MX water was ∼5 × 10^4^ CFU/mL, while the LC water, which contained a mixed microbial community, was initialized at a lower concentration of ∼5 × 10^3^ CFU/mL (Fig. 7). Analysis with a mixed measures ANOVA testing the effect of bacterial treatment (MX vs. LC), presence of larvae (present vs. absent) and day of sampling revealed significant main effects of microbiota treatment (*P* = 6.67 × 10^−9^), larval presence (*P* = 1.14 × 10^−5^), and day (*P* = 4.00 × 10^−3^), as well as two two-way interactions: microbiota treatment × larval presence (*P* = 0.020) and larval presence × day (*P* = 0.044). MX flasks had lower average bacterial loads than LC regardless of day or larval presence (*P* = 6.67 × 10^−9^; Fig. 7), and subsequent pairwise comparisons between MX and LC when data were parsed by larval presence confirmed this (larvae absent, P_MX vs. LC_ = 3.06 × 10^−6^; larvae present, P_MX vs. LC_ = 2.72 × 10^−7^). However, the difference between MX and LC was much more pronounced when larvae were absent compared to when they were present. Mean bacterial loads were 24.15-fold higher in LC compared to MX when larvae were absent but only 9.63-fold higher when larvae were present (Fig. 7; Fig. S9). The effect of larvae was also variable across days; bacterial load increased over time both in the presence and absence of larvae until day 8, at which point the absence of larvae correlated with an increase in bacterial load while presence correlated with a decrease (Fig. 7; Fig. S10).

**Fig. 7.**
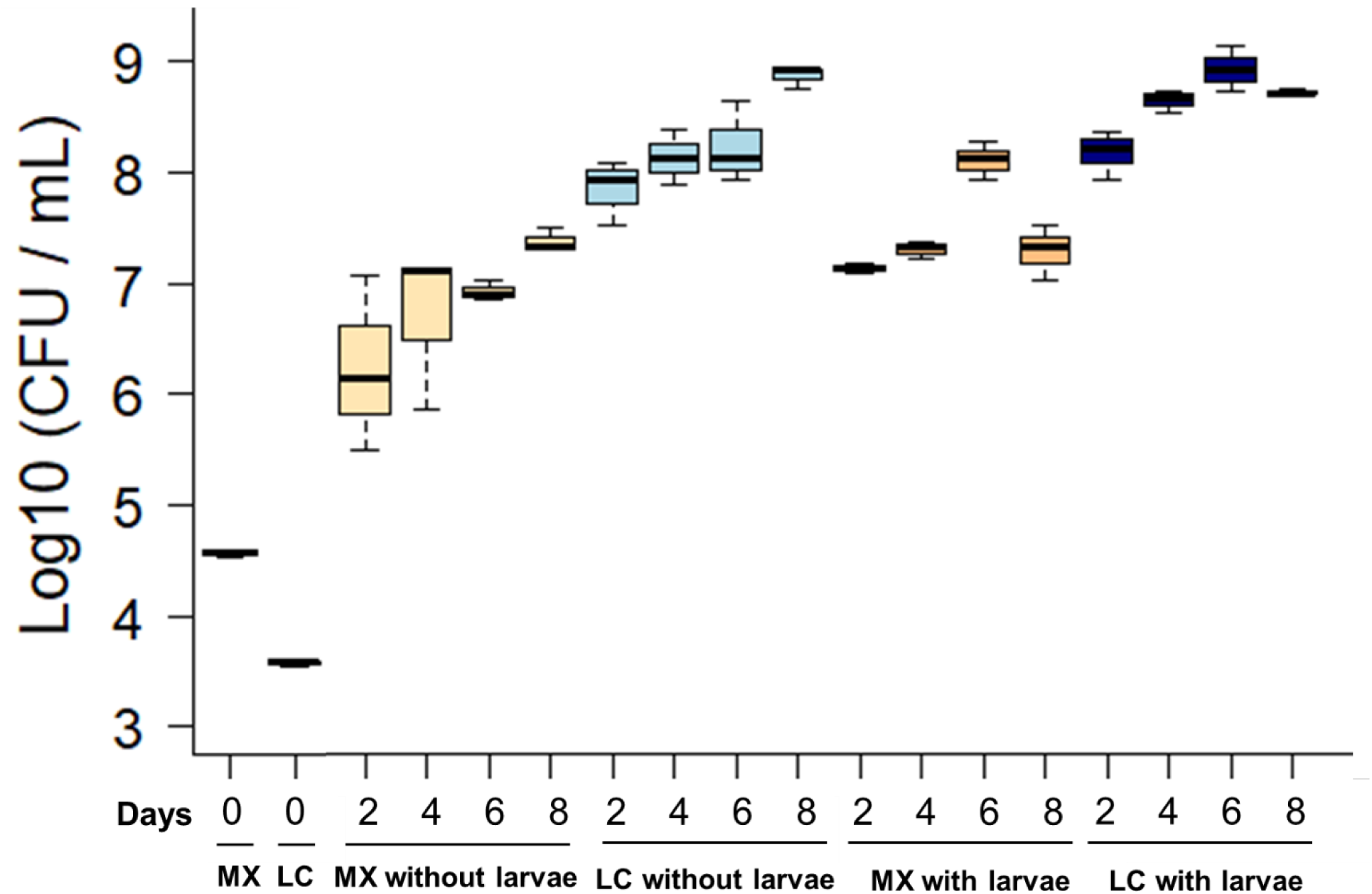
Bacterial abundance in larval rearing water is determined by microbiota treatment, larval presence, and time. We measured cultivable bacterial load in MX and LC flasks with and without larvae at 2, 4, 6, and 8 days after introducing *E. coli* S17 or the laboratory microbial community. Each box plot represents data collected from three independent replicates (i.e. flasks) for each group. Inoculation doses at day zero were calculated by culturing a dilution series from each inoculum on LB agar. Mixed measures ANOVA testing the effects of microbiota treatment, larval presence, and day on bacterial load revealed significant main effects of microbiota treatment (*P* = 6.67 × 10^−9^), larval presence, (*P* = 1.14 × 10^−5^), and day (*P* = 4.00 × 10^−3^). We found that the difference in bacterial load between MX and LC was significantly larger when larvae were absent than when they were present (microbiota treatment x larval presence, *P* = 0.020). We also found that the impact of larvae on bacterial abundance changed over time (larval presence x day, *P* = 0.044).

### Axenic treatment slows development rate, reduces male wing size and lengthens adult male longevity compared to a monoxenic *E. coli* microbiota and a laboratory microbiota

We compared AX, MX, and LC *Ae. aegypti* with regards to larval development, adult male wing length, and survival (Fig. 8A to D). In terms of larval development, AX larvae showed a significant delay in both pupation (Fig. 8A, *P*_AX vs. MX_ < 2 × 10^−16^; *P*_AX vs. LC_ < 2 × 10^−16^) and eclosion (Fig. 8B, *P*_AX vs. MX_ < 2 × 10^−16^; *P*_AX vs. LC_ < 2 × 10^−16^) compared to MX and LC larvae, but there were no significant differences in pupation or eclosion rate between the MX and LC groups (pupation rate, *P*_MX vs. LC_ = 0.088; eclosion rate, *P*_MX vs. LC_ = 0.2). AX males exhibited a significantly higher survival rate than MX males and LC males (Fig. 8C, *P*_AX vs. MX_ = 7.8 × 10^−3^; *P*_AX vs. LC_ = 1.3 × 10^−5^). A significant difference in survival was also observed between MX and LC males, with MX living longer than LC (*P*_MX vs. LC_ = 0.0412). For adult male wing length, AX males had significantly shorter wings compared to both MX and LC males (Fig. 8D, *P*_AX vs. MX_ < 1 × 10⁻^6^; *P*_AX vs. LC_ < 1 × 10^−6^). No significant difference in wing length was found between MX and LC males (*P*_MX vs. LC_ = 0.249).

**Fig. 8.**
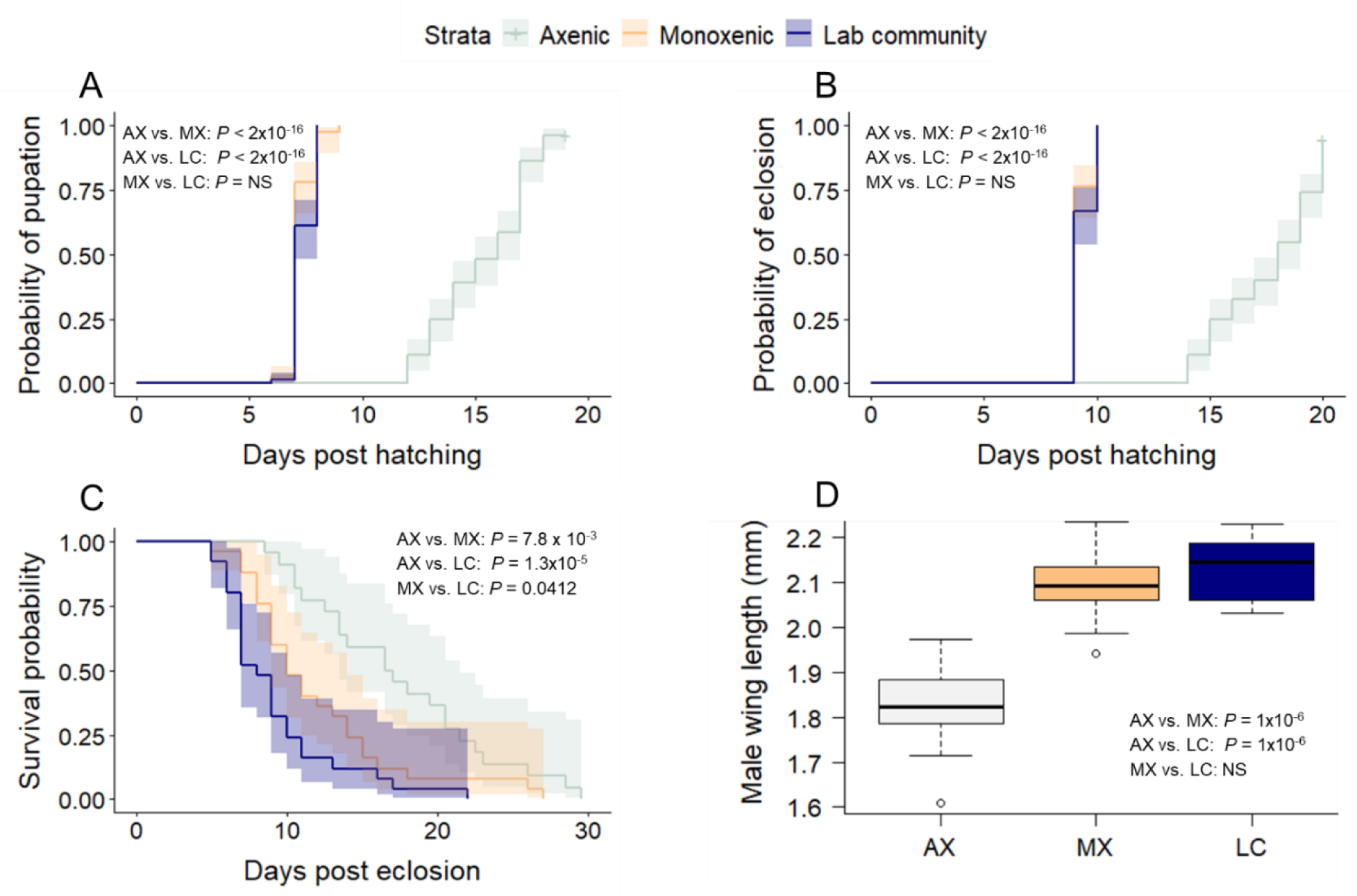
AX treatment slows larval development, reduces wing length, and enhances adult male longevity compared to MX and LC treatments. We measured pupation rate, eclosion rate, male wing length and adult male longevity for AX, MX, and LC individuals. Curves were generated using the Kaplan Meier method, and shaded areas around curves represent 95% confidence intervals. The experiment was conducted in three independent replicates for pupation and eclosion, with *n* = 24-36 mosquitoes per treatment per replicate. Survival and wing length data were collected from two other independent replicates, with *n* = 10-15 mosquitoes per treatment per replicate for survival and n = 5-12 mosquitoes per treatment per replicate for wing length. (A) Pupation rate was significantly slower for AX compared to MX (*P* < 2 × 10^−16^) and LC (*P* < 2 × 10^−16^) treatments, while no significant difference was observed between MX and LC (*P* = 0.088). (B) Similarly, eclosion rate was significantly slower in AX compared to MX (*P* < 2 × 10^−16^) and LC (*P* < 2 × 10^−16^) treatments, but no significant difference was detected between MX and LC (*P* = 0.2). (C) AX males exhibited significantly higher survival rates compared to MX (*P* = 7.8 × 10^−3^) and LC males (*P* = 1.3 × 10^−5^). Additionally, a significant difference in survival was observed between the MX and LC groups (*P* = 0.0412). (D) AX males had significantly shorter wing lengths than MX (*P* < 1 × 10⁻^6^) and LC males (*P* < 1 × 10⁻^6^), while no significant difference in wing length was observed between MX and LC groups (*P* = 0.249).

### Axenic treatment has no effect on pupation rate or adult male longevity when applied late in larval development

In a separate experiment, we sought to isolate the role of the microbiota during the adult stage specifically. We generated four treatment groups derived from MX larvae: 1) AX_Amp_ were generated by treating MX late instar larvae with ampicillin shortly prior to pupation to eliminate the microbiota; 2) AX_Amp+*E. coli*-fed_ were generated identically to AX_Amp_, then exposed to *E. coli* by feeding as adults; 3) MX were identical to previous experiments; and 4) MX*_E. coli_*_-fed_ were MX that were exposed to *E. coli* by feeding as adults. We evaluated pupation rate (Fig. S11), eclosion rate (Fig. S12), and adult male survival in all groups (Fig. 9). The absence of a microbiota in adulthood did not affect male survival, as evidenced by the comparable survival rates of AX_Amp_ males versus AX_Amp+*E. coli*-fed_ males (Fig. 9, *P* _AXAmp vs. AXAmp+*E. coli*-fed_ *=* 0.85). In addition, our findings showed that the use of ampicillin during the later larval developmental stages had no effect on pupation rate (Fig. S11, *P* = 0.595), eclosion rate (Fig. S12, *P =* 0.582) or survival of adult males (Fig. 9, *P*_AXAmp*+E. coli*-fed vs. MX*E. coli*-fed_ = 0.97). Moreover, in males that never had their microbiota removed via ampicillin treatment, ingestion of *E. coli* as an adult had no effect on male longevity (Fig. 9, *P*_MX vs. MX*E. coli*-fed_ = 0.79)

**Fig. 9.**
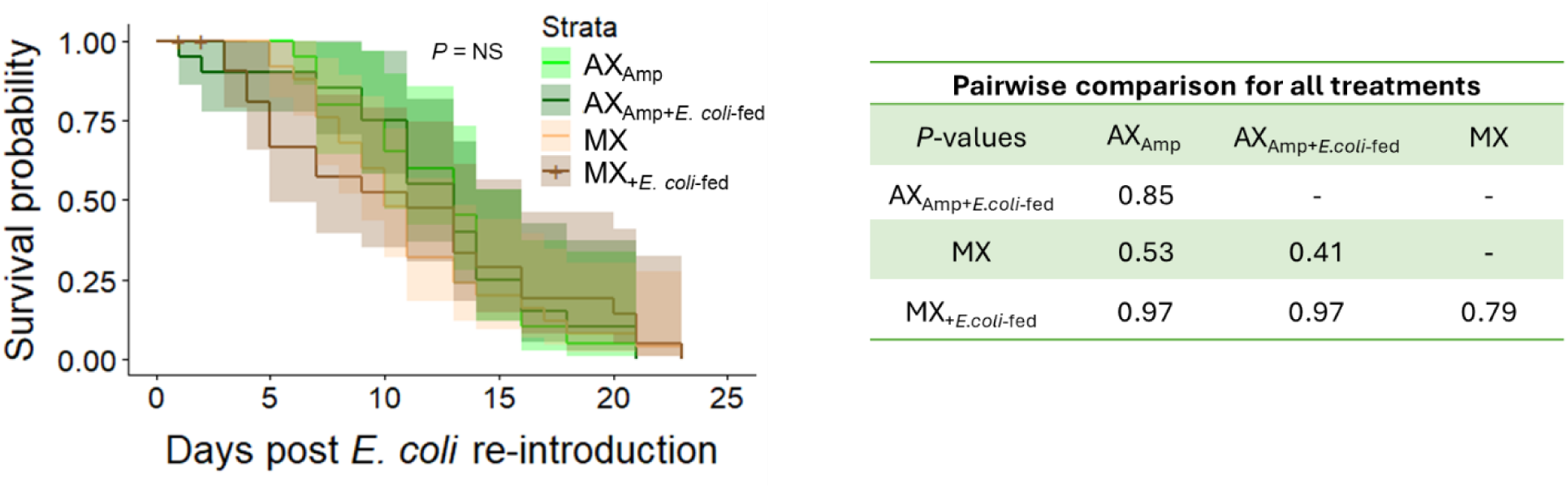
AX treatment applied late in larval development has no effect on adult male longevity. Four treatment groups were generated from MX larvae: 1) AX_Amp_ was treated with ampicillin shortly prior to pupation to eliminate the microbiota; 2) AX_Amp+*E. coli*-fed_ was generated identically to AX_Amp_, then exposed to *E. coli* by feeding as adults; 3) MX is identical to previous experiments; and 4) MX*_E. coli_*_-fed_ were MX that were also exposed to *E. coli* by feeding as adults. We evaluated survival in all groups of adult males. Curves were generated using the Kaplan Meier method, and shaded areas around curves represent 95% confidence intervals. Survival data were collected from three replicates with *n* = 9-13 mosquitoes per treatment per replicate. The absence of a microbiota in adult males did not influence their survival, as indicated by the comparable survival rates between AX_Amp_ males versus AX_Amp+*E. coli*-fed_ males (*P =* 0.85). Moreover, treatment with ampicillin did not impact adult male survival as revealed by comparable survival rates between AX_Amp*+E. coli*-fed_ vs. MX*_E. coli_*_-fed_ (*P* = 0.97), and comparison of MX and MX*_E. coli_*_-fed_ fed revealed no impact of *E. coli* ingestion on longevity among males that never had their microbiota removed via ampicillin treatment (*P* = 0.79).

## Discussion

In the current work, we show that microbiota manipulation and/or removal significantly impacts multiple life history traits in adult male *Ae. aegypti*, including lifespan, starvation resistance, and mating success. This comprehensive evaluation of the effect of the microbiota on male *Ae. aegypti* life history traits provides key foundational knowledge regarding the role of the microbiota in the biology of the yellow fever mosquito.

Our results reveal that the microbiota significantly impacts adult male lifespan. We found that adult males reared MX (i.e. inoculated with only *E. coli*) from hatching were significantly longer-lived compared to those reared LC (i.e. inoculated with a laboratory microbial community) and this lifespan extension was even more pronounced in AX (i.e. microbe-free) males. Harrison et al., (2023) reported that males reared MX as larvae and then rendered AX immediately before pupation had a prolonged lifespan compared to those that were LC as larvae and adults. Moreover, prolonged longevity in AX and MX adult females compared to LC females has been previously reported (Correa et al., 2018), as have differences in longevity between females reared MX with different bacteria (Giraud et al., 2022). Therefore, the microbiota appears to be a key determinant of adult longevity in both male and female *Ae. aegypti*.

Because it remains unclear to what extent microbiota-mediated effects on longevity are driven by the larval versus adult microbiota (e.g. Harrison et al., 2023), we conducted an experiment to incisively investigate adult-stage impacts of the microbiota on longevity. We treated late 4^th^ instar MX larvae with antibiotics and then re-colonized a subset of these individuals with *E. coli.* This generated males that differed in their adult microbiota (axenic versus monoxenic *E. coli*) but that had an identical microbiota (*E. coli* monoxenic) throughout larval development. We found no significant differences in longevity between these groups, suggesting that removal of microbiota late in larval development does not meaningfully affect adult longevity. It also suggests that the differences in longevity we observed between LC, MX, and AX individuals in Figures 3 and 8 were primarily driven by microbiota differences during the larval stages. This is consistent with findings from Romoli et al., (2024), in which the authors use an auxotrophic strain of *E. coli* to generate AX males without the use of antibiotics. They also reported no difference in lifespan between MX and AX adult males. Taken together, our full dataset suggests that the larval, but not the adult microbiota is a strong determinant of adult male lifespan.

One way in which the larval microbiota may mediate adult lifespan is via larval nutrition. Joy et al., (2010) found that adult females fed reduced larval diet amounts showed significantly increased adult lifespan compared to females fed twice the standard amount of larval diet. Given that the microbiota is an important component of the larval diet and provides necessary micronutrients (Merritt et al., 1992; Wang et al., 2021; Romoli et al., 2021), it is possible that prolonged lifespan could be driven by reduced nutritional availability in MX and/or AX larvae compared to those that develop under LC conditions. In *Drosophila melanogaster* and *Caenorhabditis elegans*, interactions between gut microbiota dysbiosis, immune system signaling, and tissue damage have been shown to impact lifespan (Clark and Walker, 2018). Such inter-system interactions remain under-investigated in mosquitoes. In light of our findings identifying a marked effect of the larval microbiota on adult male longevity, further investigation into the relationships between the microbiota, innate immunity, and gut homeostasis in mosquitoes is warranted.

The role of the microbiota on starvation resistance in mosquitoes remains largely untested (but see Giraud et al., 2022), and our findings indicate that MX males experience a significant extension of their lifespan under starvation (water only) conditions compared to LC males. Similarly to longevity, larval diet has been shown to significantly impact adult male starvation resistance. For example, survival rate of *Ae. aegypti* adult males under starvation conditions is significantly increased when larvae are fed a carbohydrate rich diet compared to a protein rich diet (Sasmita et al., 2019). Additionally, *Ae. aegypti* males reared with lower amounts of larval diet have reduced survival rates under starvation conditions compared to males fed a higher amount of larval diet (Aldersley et al., 2019). Therefore, differences in nutritional conditions between MX and LC treatments could conceivably underlie the differences in starvation resistance we observed. Intriguingly, we observed that MX males live *longer* than LC males under starvation conditions, which would not be expected if larvae are nutrient restricted (Aldersley et al., 2019). This, taken in combination with the observation that MX males have increased longevity and shorter wings, both of which *would* be expected under nutrient restriction (Ponlawat and Harrington, 2007; Joy et al., 2010; Aldersley et al., 2019), suggests microbiota manipulation impacts adult starvation resistance via a mechanism other than nutrient availability or that changes in nutrient availability induced by microbiota manipulation may not have the same effect on starvation resistance as gross manipulation of larval diet. Alternatively, starvation resistance may also vary based on the composition of the microbial community in the LC treatment. In *Drosophila melanogaster*, the microbiota has been found to dramatically impact starvation resistance (Judd et al., 2018). AX flies and flies MX for lactic acid bacterial isolates have much higher starvation resistance than flies MX for other isolates or colonized with a five-member microbial community. This difference in starvation resistance is associated with bacterial methionine metabolism genes found in the *D. melanogaster* microbiota. Increased bacterial methionine metabolism is associated with reduced starvation resistance; thus, flies harboring microbial communities enriched in bacterial methionine metabolism genes show reduced starvation resistance. Whether this is also true in mosquitoes requires further investigation, but this is one potential mechanism by which differences in bacterial community structure could induce changes in starvation resistance.

In addition to its effects on lifespan, our findings also suggest that the microbiota influences some but not all measures of mating success. When given 30 minutes to mate in a group mating non-competitive scenario, MX males were significantly more likely to inseminate females than LC males. Additionally, when given only five minutes to mate in an individual non-competitive scenario, we found that MX males had a higher probability of mating success compared to LC males, but the effect was dependent on both male and female wing length. Moreover, MX males were not more successful when directly competing with LC males for a mate, suggesting that while there are some differences in mating competency between MX and LC males, these differences do not necessarily confer a change in competitive ability. We note that we only measured wing length for the individual non-competitive scenario experiment, so it is possible that wing length could also play a role in the other mating scenarios, but we do not have the ability to assess that here. Adult body size has been shown in many studies to be relevant to mating success, but whether a particular body size is likely to confer a mating benefit is not straightforward and depends on multiple other factors. For example, small body size as a result of reduced diet provisioning results in reduced male swarming activity (Lang et al., 2018), but not necessarily a change in ability to compete for a mate (Lang et al., 2018; Aldersley et al., 2019). Additionally, size assortative mating has been reported for *Ae. aegypti*; smaller males have been reported to outcompete larger males for small, but not large mates (Callahan et al., 2018), though this has not been observed in all studies investigating the effect of male body size on mating competitiveness (Lang et al., 2018). Moreover, Cator and Zanti (2016) showed that copulation success is determined by the interaction between wing length of males and females as well as the harmonic convergence of male and female flight tone frequencies (Cator and Zanti, 2016). In our study, we also found size assortative mating; for LC males there was a positive relationship between male and female wing length in copulating pairs. This is largely consistent with the findings of Callahan et al., (2018), though we note the experiment shown in Fig. 4 did not involve male-male competition between microbiota treatments. Intriguingly, MX males deviate from the pattern in LC males; their wing length negatively correlates with that of their mates. The underlying reason for this reversal is unclear, but it presents the intriguing possibility that, in addition to having the capacity to affect male wing length, the microbiota may impact aspects of mating success that we did not measure here, such as harmonic convergence, swarming behavior, or flight capacity.

In considering the effect of the microbiota on pupation and wing length, we found that AX larvae were the slowest to pupate and had the shortest wings, while MX larvae were either slightly delayed with slightly smaller wings (Fig. 3) or no different (Fig. 8) from LC larvae depending on rearing conditions. These findings are consistent with previous studies which have shown that both AX and MX larvae successfully pupate, but the rate of AX pupation is dramatically delayed, and AX adults are significantly smaller when reared using the method we employed here (Coon et al., 2014; Correa et al., 2018; Wang et al., 2021). The delay in development is largely due to a deficit of micronutrients, especially B vitamins (Wang et al. 2021). We observed a slight delay in pupation and significantly shorter wings among the MX individuals in Fig. 3, which deviates from previous reports (Correa et al., 2018). Martinson and Strand (2021) showed that delays in pupation in MX larvae can arise under certain dietary conditions. In Fig. 3, we reared larvae at slightly higher densities than was used previously (Correa et al., 2018), and it is possible that higher larval densities may have increased larval competition for resources, thus leading to the slight delay in pupation and the reduced wing length we observed. In support of this, we found no difference in pupation rate or wing length between LC vs. MX larvae in Fig. 8, where we reared larvae under lower densities.

Finally, we analyzed bacterial abundance in larval rearing water to understand the microbial dynamics in MX vs. LC and the effects of the presence of larvae. MX larval rearing water consistently had lower bacterial loads, which may be driven by the fact that the laboratory community has a diversity of aerobic and facultative anaerobic microorganisms, some of which likely grow more rapidly in the larval rearing environment than *E. coli.* Bacterial load differences manifest by day two and remain consistent across days, suggesting that these differences are established during the earliest larval developmental stages. We also found a significant effect of larval presence on bacterial load that varied across days. At the earliest stages of larval development, flasks with larvae had higher bacterial loads than flasks without larvae. Mosquito larvae actively excrete nitrogenous waste (mainly ammonia/NH₄⁺) (Donini & O’Donnell, 2005; Weihrauch & Allen, 2018; Durant & Donini, 2019) into the rearing water and these inputs could stimulate microbial growth and/or shifts in bacterial community composition. Accordingly, larvae actively shape microbial abundance by enriching the environment with organic inputs. The presence of *Aedes triseriatus* larvae in microcosms has been shown to increase the abundance of aerobic and facultative anaerobic bacteria in water samples and correlates with significant shifts in the presence of organic compounds in the water column (Kaufman et al., 1999). By day eight, when most of the larvae had pupated, flasks with larvae/pupae had lower bacterial loads than those without larvae/pupae. This may reflect cessation of larval waste excretion and subsequent reduction in nutrient availability for microbial growth. Or, not mutually exclusively, it may be related to the fact that bacterial populations in the mosquito midgut decline during metamorphosis from larva to pupa and adult (Walker & Romoser, 1987; Moll et al., 2001). Although these studies primarily describe internal gut microbiota, it is possible that reductions in bacterial load associated with molting to the pupal stage indirectly influence microbial dynamics in the rearing environment. However, the mechanisms linking mosquito development to changes in water bacterial abundance remain unclear and need further investigation. Overall, these results showed that bacterial treatment, larval presence, and time independently contributed to increasing bacterial abundance, with larvae playing a dynamic role in shaping microbial communities during their development. These findings emphasize that the larval microbiota interaction is not static but associated with larval development over time.

Our findings underscore the intricate interactions between *Aedes aegypti* male mosquitoes and the microbes in their larval environment, shedding light on how the microbiota influences key male life history traits that impact male mosquito fitness. The presence or absence of a microbiota affects multiple male life history traits, including development, body size, longevity, and mating success, highlighting the complex role of microbial communities in mosquito biology. These findings provide valuable insights into the impact of the microbiota on mosquito biology and have important implications for vector control strategies that utilize mass rearing.

## Supporting information

File S2 - R code and output

File S1 - All raw data

## Acknowledgements

We thank Laura Harrington, Cornell University, for providing Thai strain *Ae. aegypti* and George Dimopoulos, Johns Hopkins Bloomberg School of Public Health, for providing *E. coli* S17 pPROBE-mCherry.

