## Supplementary material for "The microbiota impacts life history traits and mating success in male *Aedes aegypti* mosquitoes": File S2 - R code and output

### Experiment 1: Comparing development time and adult male life history/mating traits between MX and LC individuals.

##### Setup: Load R packages and set plot margins

library(tidyverse)
library(survival)
library(survminer)
library(interactions)
library(arm)
library(stats)
library(car)
library(DHARMa)
library(ggpubr)
library(rstatix)
library(lavaan)
library(lavaanPlot)
library(GGally)
par(mar=c(10.2,8.2,8.2,4.2))

#### Fig (3A): Experiment 1, Pupation Analysis (MX vs LC)

##### **R Code**

```
### Load and format data
mxlc_pup <- read.table("Fig3A_MX_LC_Pupation.csv", sep=",", header=TRUE)
mxlc_pup$treatment <- factor(mxlc_pup$treatment, levels=c("mx", "lc")) #set reference level
mxlc_pup$rep <- as.factor(mxlc_pup$rep)

### Analyses
#fit cox proportional hazards model:
cox1 <- coxph(Surv(daysposthatching, status, type=c('right'))~treatment+rep, data=mxlc_pup)
cox.zph(cox1, global=T) #coxph assumptions violated for treatment and rep
#perform logrank test to assess overall effect of treatment on pupation:
survdiff(Surv(daysposthatching, status)~treatment, data=mxlc_pup) #treatment p=<2e-16
#perform logrank test to assess effect of treatment on pupation within each replicate:
mxlc_pup_rep1 <- mxlc_pup %>% dplyr::filter(rep=="one")
mxlc_pup_rep2 <- mxlc_pup %>% dplyr::filter(rep=="two")
mxlc_pup_rep3 <- mxlc_pup %>% dplyr::filter(rep=="three")
mxlc_pup_rep4 <- mxlc_pup %>% dplyr::filter(rep=="four")
survdiff(Surv(daysposthatching, status)~treatment, data=mxlc_pup_rep1) #treatment p=2e-15
survdiff(Surv(daysposthatching, status)~treatment, data=mxlc_pup_rep2) #treatment p=<2e-16
survdiff(Surv(daysposthatching, status)~treatment, data=mxlc_pup_rep3) #treatment p=7e-07
survdiff(Surv(daysposthatching, status)~treatment, data=mxlc_pup_rep4) #treatment p=<2e-16

### Figures
##create fig3a
km.mxlc_pup <- survfit(Surv(daysposthatching, status)~treatment, data=mxlc_pup) #fit survival object
fig3a <- ggsurvplot(
  km.mxlc_pup,
  palette=c("#FFBF80", "navy"),
  legend.labs =c("Monoxenic", "Lab Community"),
  conf.int=TRUE,
  fun="event",
  ylab="Probability of pupation",
  xlab="Days post hatching",
  break.time.by=2,
  font.x=c(20),
  font.y=c(20),
  font.tickslab=c(16))
```

##### **Analysis Outputs**

```
survdiff(Surv(daysposthatching, status)~treatment, data=mxlc_pup)
## Call:
## survdiff(formula = Surv(daysposthatching, status) ~ treatment,
##     data = mxlc_pup)
##
##                N Observed Expected (O-E)^2/E (O-E)^2/V
## treatment=mx 275      258      365      31.3       218
## treatment=lc 250      250      143      79.7       218
##
##  Chisq= 218  on 1 degrees of freedom, p= <2e-16
survdiff(Surv(daysposthatching, status)~treatment, data=mxlc_pup_rep1) # replicate 1
## Call:
## survdiff(formula = Surv(daysposthatching, status) ~ treatment,
##     data = mxlc_pup_rep1)
##
##               N Observed Expected (O-E)^2/E (O-E)^2/V
## treatment=mx 50       43       67      8.61      63.3
## treatment=lc 50       50       26     22.22      63.3
##
##  Chisq= 63.3  on 1 degrees of freedom, p= 2e-15
survdiff(Surv(daysposthatching, status)~treatment, data=mxlc_pup_rep2) # replicate 2
## Call:
## survdiff(formula = Surv(daysposthatching, status) ~ treatment,
##     data = mxlc_pup_rep2)
##
##               N Observed Expected (O-E)^2/E (O-E)^2/V
## treatment=mx 75       72    106.6      11.2      86.6
## treatment=lc 75       75     40.4      29.7      86.6
##
##  Chisq= 86.6  on 1 degrees of freedom, p= <2e-16
survdiff(Surv(daysposthatching, status)~treatment, data=mxlc_pup_rep3) # replicate 3
## Call:
## survdiff(formula = Surv(daysposthatching, status) ~ treatment,
##     data = mxlc_pup_rep3)
##
##               N Observed Expected (O-E)^2/E (O-E)^2/V
## treatment=mx 75       75     90.6      2.68      24.7
## treatment=lc 50       50     34.4      7.05      24.7
##
##  Chisq= 24.7  on 1 degrees of freedom, p= 7e-07
survdiff(Surv(daysposthatching, status)~treatment, data=mxlc_pup_rep4) # replicate 4
## Call:
## survdiff(formula = Surv(daysposthatching, status) ~ treatment,
##     data = mxlc_pup_rep4)
##
##               N Observed Expected (O-E)^2/E (O-E)^2/V
## treatment=mx 75       68      103      11.9      86.2
## treatment=lc 75       75       40      30.7      86.2
##
##  Chisq= 86.2  on 1 degrees of freedom, p= <2e-16
```

#### Fig (3B): Experiment 1, Eclosion Analysis (MX vs LC)

##### **R Code**

```
### Load and format data
mxlc_eclo <- read.table("Fig3B_MX_LC_Eclosion.csv", sep=",", header=TRUE)
mxlc_eclo$treatment <- factor(mxlc_eclo$treatment, levels=c("mx", "lc")) #set reference level
mxlc_eclo$rep <- as.factor(mxlc_eclo$rep)

### Analyses
#fit cox proportional hazards model:
cox2 <- coxph(Surv(daysposthatching, status, type=c('right'))~treatment+rep, data=mxlc_eclo)
cox.zph(cox2, global=T) #coxph assumptions violated for treatment and rep
#perform logrank test to assess overall effect of treatment on eclosion:
survdiff(Surv(daysposthatching, status)~treatment, data=mxlc_eclo) #treatment p=<2e-16
#perform logrank test to assess effect of treatment on eclosion within each replicate:
mxlc_eclo_rep1 <- mxlc_eclo %>% dplyr::filter(rep=="one")
mxlc_eclo_rep2 <- mxlc_eclo %>% dplyr::filter(rep=="two")
mxlc_eclo_rep3 <- mxlc_eclo %>% dplyr::filter(rep=="three")
mxlc_eclo_rep4 <- mxlc_eclo %>% dplyr::filter(rep=="four")
survdiff(Surv(daysposthatching, status)~treatment, data=mxlc_eclo_rep1) #treatment p=1e-10
survdiff(Surv(daysposthatching, status)~treatment, data=mxlc_eclo_rep2) #treatment p=<2e-16
survdiff(Surv(daysposthatching, status)~treatment, data=mxlc_eclo_rep3) #treatment p=9e-06
survdiff(Surv(daysposthatching, status)~treatment, data=mxlc_eclo_rep4) #treatment p=4e-08

### Figures
#create fig3b
km.mxlc_eclo <- survfit(Surv(daysposthatching,status)~treatment, data=mxlc_eclo) #fit survival object
fig3b <- ggsurvplot(
  km.mxlc_eclo,
  palette=c("#FFBF80", "navy"),
  legend.labs =c("Monoxenic", "Lab Community"),
  font.legend=c(13),
  fun="event",
  conf.int=TRUE,
  ylab="Probability of eclosion",
  xlab="Days post hatching",
  font.x=c(20),
  font.y=c(20),
  font.tickslab=c(16))
```

##### **Analysis Outputs**

```
survdiff(Surv(daysposthatching, status)~treatment, data=mxlc_eclo)
## Call:
## survdiff(formula = Surv(daysposthatching, status) ~ treatment,
##     data = mxlc_eclo)
##
##                N Observed Expected (O-E)^2/E (O-E)^2/V
## treatment=mx 275      250      338      23.1       134
## treatment=lc 249      239      151      51.8       134
##
##  Chisq= 134  on 1 degrees of freedom, p= <2e-16
survdiff(Surv(daysposthatching, status)~treatment, data=mxlc_eclo_rep1) # replicate 1
## Call:
## survdiff(formula = Surv(daysposthatching, status) ~ treatment,
##     data = mxlc_eclo_rep1)
##
##               N Observed Expected (O-E)^2/E (O-E)^2/V
## treatment=mx 50       37     57.1      7.05      41.1
## treatment=lc 50       47     26.9     14.93      41.1
##
##  Chisq= 41.1  on 1 degrees of freedom, p= 1e-10
survdiff(Surv(daysposthatching, status)~treatment, data=mxlc_eclo_rep2) # replicate 2
## Call:
## survdiff(formula = Surv(daysposthatching, status) ~ treatment,
##     data = mxlc_eclo_rep2)
##
##               N Observed Expected (O-E)^2/E (O-E)^2/V
## treatment=mx 75       73    106.6      10.6      78.5
## treatment=lc 74       74     40.4      28.0      78.5
##
##  Chisq= 78.5  on 1 degrees of freedom, p= <2e-16
survdiff(Surv(daysposthatching, status)~treatment, data=mxlc_eclo_rep3) # replicate 3
## Call:
## survdiff(formula = Surv(daysposthatching, status) ~ treatment,
##     data = mxlc_eclo_rep3)
##
##               N Observed Expected (O-E)^2/E (O-E)^2/V
## treatment=mx 75       75     89.8      2.44      19.8
## treatment=lc 50       50     35.2      6.22      19.8
##
##  Chisq= 19.8  on 1 degrees of freedom, p= 9e-06
survdiff(Surv(daysposthatching, status)~treatment, data=mxlc_eclo_rep4) # replicate 4
## Call:
## survdiff(formula = Surv(daysposthatching, status) ~ treatment,
##     data = mxlc_eclo_rep4)
##
##               N Observed Expected (O-E)^2/E (O-E)^2/V
## treatment=mx 75       65     88.2      6.09      30.2
## treatment=lc 75       68     44.8     11.99      30.2
##
##  Chisq= 30.2  on 1 degrees of freedom, p= 4e-08
```

#### Fig (3C): Experiment 1, Survival Analysis in Sugar-Fed (MX vs LC)

##### **R Code**

```
### Load and format data
mxlc_death <- read.table("Fig3C_MX_LC_Survival_SugarFed.csv", sep=",", header=TRUE)
mxlc_death$treatment <- factor(mxlc_death$treatment, levels=c("mx", "lc")) #set reference level
mxlc_death$rep <- as.factor(mxlc_death$rep)

### Analyses
#fit cox proportional hazards model:
cox3 <- coxph(Surv(daysposteclosion, status, type=c('right'))~treatment+rep, data=mxlc_death)
cox.zph(cox3, global=T) #coxph assumptions met
summary(cox3)
anova(cox3) #treatment p=4.253e-07

### Figures
#create fig3c
km.mxlc_death <- survfit(Surv(daysposteclosion,status)~treatment, data=mxlc_death) #fit survival object
fig3c <- ggsurvplot(
  km.mxlc_death,
  conf.int=TRUE,
  ylab=" Male adult survival probability\n(10% sucrose)",
  xlab="Days post eclosion",
  font.x=c(20),
  font.y=c(20),
  font.legend=c(13),
  font.tickslab=c(16),
  palette=c("#FFBF80", "navy"),
  legend.labs =c("Monoxenic", "Lab Community"))
```

##### **Analysis Outputs**

```
cox3 <- coxph(Surv(daysposteclosion, status, type=c('right'))~treatment+rep, data=mxlc_death)
anova(cox3)
## Analysis of Deviance Table
##  Cox model: response is Surv(daysposteclosion, status, type = c("right"))
## Terms added sequentially (first to last)
##
##            loglik  Chisq Df Pr(>|Chi|)
## NULL      -296.92
## treatment -284.13 25.576  1  4.253e-07 ***
## rep       -273.60 21.063  2  2.668e-05 ***
## ---
## Signif. codes:  0 '***' 0.001 '**' 0.01 '*' 0.05 '.' 0.1 ' ' 1
```

#### Fig (3D): Experiment 1, Survival Analysis in Water-Fed (MX vs LC)

##### **R Code**

```
### Load and format data
mxlc_death2 <- read.table("Fig3D_MX_LC_Survival_WaterFed.csv", sep=",", header=TRUE)
mxlc_death2$treatment <- factor(mxlc_death2$treatment, levels=c("mx", "lc")) #set reference level
mxlc_death2$rep <- as.factor(mxlc_death2$rep)

### Analyses
#fit cox proportional hazards model:
cox4 <- coxph(Surv(daysposteclosion, status, type=c('right'))~treatment+rep, data=mxlc_death2)
cox.zph(cox4, global=T) #coxph assumptions violated globally (p=0.048) and for rep (p=0.020)
#because p-values>0.01, fit cox models with and without inclusion of replicate predictor to evaluate effect of treatment; additionally, evaluate effect of treatment within each individual replicate using logrank test
#fit cox proportional hazards model without rep:
cox4b <- coxph(Surv(daysposteclosion, status, type=c('right'))~treatment, data=mxlc_death2)
cox.zph(cox4b, global=T) #coxph assumptions met
anova(cox4) #treatment: p=0.01
anova(cox4b) #treatment: p=0.01
#perform logrank test to assess effect of treatment on pupation within each replicate:
mxlc_death2_rep1 <- mxlc_death2 %>% dplyr::filter(rep=="one")
mxlc_death2_rep2 <- mxlc_death2 %>% dplyr::filter(rep=="two")
mxlc_death2_rep3 <- mxlc_death2 %>% dplyr::filter(rep=="three")
survdiff(Surv(daysposteclosion, status)~treatment, data=mxlc_death2_rep1) #treatment: p=1e-04
survdiff(Surv(daysposteclosion, status)~treatment, data=mxlc_death2_rep2) #treatment: p=0.8
survdiff(Surv(daysposteclosion, status)~treatment, data=mxlc_death2_rep3) #treatment: p=0.3

### Figures
#create fig3d
km.mxlc_death2 <- survfit(Surv(daysposteclosion,status)~treatment, data=mxlc_death2) #fit survival object
fig3d <- ggsurvplot(
  km.mxlc_death2,
  conf.int=TRUE,
  ylab="Male adult survival probability \n (Water only)",
  xlab="Days post eclosion",
  font.x=c(20),
  font.y=c(20),
  font.tickslab=c(16),
  break.time.by=2,
  palette=c("#FFBF80", "navy"),
  legend.labs =c("Monoxenic", "Lab Community"),
  font.legend=c(13))
```

##### **Analysis Outputs**

```
cox4 <- coxph(Surv(daysposteclosion, status, type=c('right'))~treatment+rep, data=mxlc_death2)
anova(cox4)
## Analysis of Deviance Table
##  Cox model: response is Surv(daysposteclosion, status, type = c("right"))
## Terms added sequentially (first to last)
##
##            loglik  Chisq Df Pr(>|Chi|)
## NULL      -318.15
## treatment -314.99 6.3274  1    0.01189 *
## rep       -312.79 4.3883  2    0.11145
## ---
## Signif. codes:  0 '***' 0.001 '**' 0.01 '*' 0.05 '.' 0.1 ' ' 1
cox4b <- coxph(Surv(daysposteclosion, status, type=c('right'))~treatment, data=mxlc_death2)
anova(cox4b)
## Analysis of Deviance Table
##  Cox model: response is Surv(daysposteclosion, status, type = c("right"))
## Terms added sequentially (first to last)
##
##            loglik  Chisq Df Pr(>|Chi|)
## NULL      -318.15
## treatment -314.99 6.3274  1    0.01189 *
## ---
## Signif. codes:  0 '***' 0.001 '**' 0.01 '*' 0.05 '.' 0.1 ' ' 1
survdiff(Surv(daysposteclosion, status)~treatment, data=mxlc_death2_rep1) # replicate 1
## Call:
## survdiff(formula = Surv(daysposteclosion, status) ~ treatment,
##     data = mxlc_death2_rep1)
##
##               N Observed Expected (O-E)^2/E (O-E)^2/V
## treatment=mx 15       15    22.09      2.27      14.6
## treatment=lc 15       15     7.91      6.35      14.6
##
##  Chisq= 14.6  on 1 degrees of freedom, p= 1e-04
survdiff(Surv(daysposteclosion, status)~treatment, data=mxlc_death2_rep2) # replicate 2
## Call:
## survdiff(formula = Surv(daysposteclosion, status) ~ treatment,
##     data = mxlc_death2_rep2)
##
##               N Observed Expected (O-E)^2/E (O-E)^2/V
## treatment=mx 15       15     15.5    0.0168     0.061
## treatment=lc 15       15     14.5    0.0180     0.061
##
##  Chisq= 0.1  on 1 degrees of freedom, p= 0.8
survdiff(Surv(daysposteclosion, status)~treatment, data=mxlc_death2_rep3) # replicate 3
## Call:
## survdiff(formula = Surv(daysposteclosion, status) ~ treatment,
##     data = mxlc_death2_rep3)
##
##               N Observed Expected (O-E)^2/E (O-E)^2/V
## treatment=mx 15       15     17.3     0.311      1.22
## treatment=lc 15       15     12.7     0.425      1.22
##
##  Chisq= 1.2  on 1 degrees of freedom, p= 0.3
```

#### Fig (4A): Experiment 1, Wing length (MX vs LC)

##### **R Code**

```
### Load and format data
mxlc_maleWL <- read.table("Fig4_S678_MX_LC_Wings_Cop_Inse.csv", sep=",", header=TRUE)
mxlc_maleWL$treatment <- factor(mxlc_maleWL$treatment, levels=c("mx", "lc")) #set reference level
mxlc_maleWL$replicate <- factor(mxlc_maleWL$replicate, levels=c("one", "two", "three", "four"))

### Analyses
#assess effect of treatment on winglength with all replicates combined:
#set contrasts for type III sums of squares
contrasts(mxlc_maleWL$treatment) <- contr.sum
contrasts(mxlc_maleWL$replicate) <- contr.sum
y <- lm(malewinglength~treatment+replicate, data=mxlc_maleWL)
#check model fit
plot(y) #fit is adequate
summary(y)
#run type 3 SS anova
Anova(y, type="III", test.statistic="F") #treatment p=0.0001495

### Figures
#create fig4a
fig4a <- boxplot(
  malewinglength~treatment, data=mxlc_maleWL,
  col=c("#FFBF80", "navy"),
  cex.lab=1.5,
  cex.axis=1.1,
  ylab="Male wing length (mm)",
  names=c("MX", "LC"),  # Change x-axis labels
  ylim=c(1.6, 2.2),
  xlab="",
  las=1)  # Rotate y-axis numbers to vertical
```

##### **Analysis Outputs**

```
y <- lm(malewinglength~treatment+replicate, data=mxlc_maleWL)
Anova(y, type="III", test.statistic="F")
## Anova Table (Type III tests)
##
## Response: malewinglength
##              Sum Sq Df    F value    Pr(>F)
## (Intercept) 298.665  1 25696.5048 < 2.2e-16 ***
## treatment     0.186  1    15.9877 0.0001495 ***
## replicate     0.164  3     4.6925 0.0046962 **
## Residuals     0.860 74
## ---
## Signif. codes:  0 '***' 0.001 '**' 0.01 '*' 0.05 '.' 0.1 ' ' 1
```

#### Fig (4B-D, S8): Experiment 1, Non-competitive mating (Individual)

##### **R Code**

```
### Load and format data
mxlc_IndivMating <- read.table("Fig4_S678_MX_LC_Wings_Cop_Inse.csv", sep=",", header=TRUE)
mxlc_IndivMating$treatment <- factor(mxlc_IndivMating$treatment, levels=c("mx", "lc")) #set reference level
mxlc_IndivMating$replicate <- factor(mxlc_IndivMating$replicate, levels=c("one", "two", "three", "four"))
is.ordered(mxlc_IndivMating$treatment) #confirm treatment is an unordered factor
is.ordered(mxlc_IndivMating$replicate) #confirm replicate is an unordered factor
```

###### TIME TO COPULA FORMATION: GLM

```
###Analyses
#fit general linear model:
y1 <- glm(timetocopula~treatment*femwinglength*malewinglength+replicate, family="gaussian", data=mxlc_IndivMating)
summary(y1)
#check model fit using DHARMa
plot(simulateResiduals(fittedModel=y1)) #quantile deviations detected, suggesting inadequate fit
#fit general linear model without 3-way interaction:
y1 <- glm(timetocopula~treatment*femwinglength+treatment*malewinglength+femwinglength*malewinglength+replicate, family="gaussian", data=mxlc_IndivMating)
summary(y1)
#check model fit using DHARMa
plot(simulateResiduals(fittedModel=y1)) #the fit is adequate
#run type 3 SS anova
Anova(y1, type="III", test.statistic="F") #no significant predictors

### Figures
#create fig4b
fig4b <- boxplot(
  timetocopula~treatment,
  data=mxlc_IndivMating,
  col=c("#FFBF80", "navy"),  # Ensure colors match treatment order
  cex.lab=1.5,
  cex.axis=1.1,
  ylab="Time to copula formation (s)",
  names=c("MX", "LC"),  # Change x-axis labels
  xlab="",
  ylim=c(15, 305),
  las=1)  # Rotate y-axis numbers to vertical
```

###### COPULATION SUCCESS: GLM

```
### Analyses
#fit binomial model to test effect of treatment, male wing length and female wing length on copula #formation
y2 <- glm(copulaformation~treatment*femwinglength*malewinglength+replicate, family="binomial", data=mxlc_IndivMating)
summary(y2)
#check model fit using DHARMa
plot(simulateResiduals(fittedModel=y2)) #the fit is adequate
#run type 3 SS anova
Anova(y2, type="III", test.statistic="LR") #treatment p=0.033, treatment*femwinglength*malewinglength p=0.034

### Figures
#create figS8 interaction plot to visualize the 3-way interaction:
figS8<- interact_plot(y2, pred=femwinglength, modx=treatment, mod2=malewinglength, interval=T, int.width=0.8,
                      colors= c("#FFBF80","navy"), x.label="Female wing length", y.label="Probability of Copulation" )
#create fig4d (parsing data by whether copulation was successful):
copulaformation.labs <- c("Unsuccessful copulation", "Successful copulation")
names(copulaformation.labs) <- c("0", "1")
fig4d <- ggplot(
  aes(x=femwinglength,
      y=malewinglength,
      color=treatment,
      fill=treatment),
  data=mxlc_IndivMating) +
  geom_smooth(method="lm", se=TRUE, size=1.5) +
  geom_point(size=1.5, shape=21, color="black") +
  scale_color_manual(values=c("#FFBF80", "#000080"), name="Treatment", labels=c("MX", "LC")) +
  scale_fill_manual(values=c("#FFBF80", "#000080"), name="Treatment", labels=c("MX", "LC")) +
  facet_grid(~copulaformation, labeller=labeller(copulaformation=copulaformation.labs)) +
  ylab("Male wing length (mm)") +
  xlab("Female wing length (mm)") +
  theme_pubr() +
  theme(axis.text.x=element_text(size=12),
        axis.text.y=element_text(size=12),
        axis.title.x=element_text(size=16),
        axis.title.y=element_text(size=16),
        panel.border=element_rect(),
        strip.background=element_rect())
#create fig4c
#the y2 type III SS anova assesses main effects in presence of interactions
copulaformation.predicted <- predict(y2, mxlc_IndivMating, type="response") #compute the predicted values
mxlc_IndivMating$copulaformation.predicted <- copulaformation.predicted #add it back into the dataframe
fig4c <- boxplot(
  copulaformation.predicted~treatment, data=mxlc_IndivMating,
  col=c("#FFBF80", "navy"),  # Ensure colors match treatment order
  cex.lab=1.3,
  cex.axis=1.1,
  ylab="Predicted probability of copulation success",
  names=c("MX", "LC"),  # Change x-axis labels
  xlab="",
  ylim=c(0,1),
  las=1)  # Rotate y-axis numbers to vertical
```

##### **Analysis Outputs**

###### TIME TO COPULA FORMATION: GLM

```
y1 <- glm(timetocopula~treatment*femwinglength+treatment*malewinglength+femwinglength*malewinglength+replicate, family="gaussian", data=mxlc_IndivMating)
Anova(y1, type="III", test.statistic="F")
## Analysis of Deviance Table (Type III tests)
##
## Response: timetocopula
## Error estimate based on Pearson residuals
##
##                              Sum Sq Df F values Pr(>F)
## treatment                      4677  1   0.8736 0.3553
## femwinglength                 13316  1   2.4873 0.1223
## malewinglength                10982  1   2.0514 0.1595
## replicate                     21676  3   1.3496 0.2713
## treatment:femwinglength        9014  1   1.6838 0.2015
## treatment:malewinglength       1033  1   0.1929 0.6628
## femwinglength:malewinglength  10411  1   1.9447 0.1705
## Residuals                    224845 42
```

###### COPULATION SUCCESS: GLM

```
y2 <- glm(copulaformation~treatment*femwinglength*malewinglength+replicate, family="binomial", data=mxlc_IndivMating)
Anova(y2, type="III", test.statistic="LR")
## Analysis of Deviance Table (Type III tests)
##
## Response: copulaformation
##                                        LR Chisq Df Pr(>Chisq)
## treatment                                4.5326  1    0.03326 *
## femwinglength                            3.0380  1    0.08134 .
## malewinglength                           3.0206  1    0.08221 .
## replicate                               10.1695  3    0.01718 *
## treatment:femwinglength                  4.4974  1    0.03395 *
## treatment:malewinglength                 4.5340  1    0.03323 *
## femwinglength:malewinglength             3.0062  1    0.08295 .
## treatment:femwinglength:malewinglength   4.4955  1    0.03398 *
## ---
## Signif. codes:  0 '***' 0.001 '**' 0.01 '*' 0.05 '.' 0.1 ' ' 1
```

#### Fig (S6 and S7) and Table (S1 and S2): Experiment 1, Non-competitive mating (Individual) - Path Analysis

##### **R Code**

```
# https://lavaan.ugent.be/tutorial/
# load and format data
mxlc_IndivMating <- read.table("Fig4_S678_MX_LC_Wings_Cop_Inse.csv", sep=",", header=TRUE)
mxlc_IndivMating$treatment <- factor(mxlc_IndivMating$treatment, levels=c("mx", "lc"))
mxlc_IndivMating$replicate <- factor(mxlc_IndivMating$replicate, levels=c("one", "two", "three", "four"))
mxlc_IndivMating$treatment <- factor(mxlc_IndivMating$treatment, levels=c("mx", "lc"))
mxlc_IndivMating$trmt <- mxlc_IndivMating$treatment
mxlc_IndivMating <- mxlc_IndivMating[,c(1,2,10,3,4,5,6,7,8,9)] #reorder columns
# graph data to visually assess relationships & check for correlations
plot(mxlc_IndivMating[, 3:8])
cp <- ggpairs(mxlc_IndivMating[, c(3:6,8)], upper=list(continuous=wrap("cor", size=5, color="black")))
cp + theme(strip.text.x=element_text(size=18), strip.text.y=element_text(size=10))
hist(mxlc_IndivMating$malewinglength)
plot(mxlc_IndivMating$malewinglength, pch=19)
boxplot(mxlc_IndivMating$malewinglength ~ mxlc_IndivMating$treatment)
hist(mxlc_IndivMating$femwinglength)
plot(mxlc_IndivMating$femwinglength, pch=19)
boxplot(mxlc_IndivMating$femwinglength ~ mxlc_IndivMating$treatment)
hist(mxlc_IndivMating$timetocopula)
plot(mxlc_IndivMating$timetocopula, pch=19)
plot(mxlc_IndivMating$timetocopula ~ mxlc_IndivMating$malewinglength, pch=19)
boxplot(mxlc_IndivMating$timetocopula ~ mxlc_IndivMating$treatment)
mxlc_IndivMating.cp <- mxlc_IndivMating[!is.na(mxlc_IndivMating$timetocopula), ]
mxlc_IndivMating.cp$timetocopula.scale <- scale(mxlc_IndivMating.cp$timetocopula, center=TRUE, scale=TRUE)
str(mxlc_IndivMating.cp)
mxlc_IndivMating$copulaformation <- ordered(mxlc_IndivMating$copulaformation) #need to convert binary response variable to ordered factor for these analysis
```

###### TIME TO COPULA FORMATION: PATH ANALYSIS

```
## specify the model
mod1 <- "
# direct effects
copulaformation ~ b * malewinglength + c * treatment
# mediator
malewinglength ~ a * treatment
# indirect effects
trt.malew := a * b
# total effects
total := a * b + c
"
## fit the model
fit_mod1 <- sem(mod1, #bootstrapping will produce a different output each time the model is fit
                data=mxlc_IndivMating,
                ordered=c("copulaformation"),
                model.type="cfa",
                estimator="DWLS",
                se="bootstrap",
                bootstrap=1000)
# view summary statistics (Table S1)
summary(fit_mod1,
        fit.measures=TRUE,
        standardized=TRUE,
        rsquare=TRUE,
        ci=TRUE)
parameterestimates(fit_mod1)
# Create SEM diagram (Fig. S6)
lavaanPlot(fit_mod1, coefs=TRUE, covs=TRUE, stars="regress")
```

###### COPULATION SUCCESS: PATH ANALYSIS

```
## specify the model
mod2 = "
# direct effects
timetocopula.scale ~ b * malewinglength + c * treatment
# mediator
malewinglength ~ a * treatment
# indirect effects
trt.malew := a * b
# total effects
total := a * b + c
"
## fit model
fit_mod2 <- sem(mod2, #bootstrapping will produce a different output each time the model is fit
                data=mxlc_IndivMating.cp,
                model.type="cfa",
                estimator="ML",
                se="bootstrap",
                bootstrap=1000)
# view summary statistics (Table S2)
summary(fit_mod2,
        fit.measures=TRUE,
        standardized=TRUE,
        rsquare=TRUE,
        ci=TRUE)
parameterestimates(fit_mod2)
# Create SEM diagram (Fig. S7)
lavaanPlot(fit_mod2, coefs=TRUE, covs=TRUE, stars="regress")
```

#### Fig (5A and B): Experiment 1, Non-competitive mating (Groups)

##### **R Code**

```
### Load and format data
mxlc_GroupMating <- read.table("Fig5_MX_LC_GrpMtng_Binomial.csv", sep=",", header=TRUE)
mxlc_GroupMating$treatment <- factor(mxlc_GroupMating$treatment, levels=c("mx", "lc")) #set reference level
mxlc_GroupMating$replicate <- as.factor(mxlc_GroupMating$replicate)
mxlc_GroupMating$minutes <- factor(mxlc_GroupMating$minutes, ordered=TRUE)
is.ordered(mxlc_GroupMating$treatment) #confirm treatment is an unordered factor
is.ordered(mxlc_GroupMating$replicate) #confirm replicate is an unordered factor
is.ordered(mxlc_GroupMating$minutes) #confirm minutes is an ordered factor

### Analyses
#fit glm
y13 <- glm(spermtransfer~treatment*minutes+replicate, family="binomial", data=mxlc_GroupMating)
summary(y13)
#check model fit using DHARMa
plot(simulateResiduals(fittedModel=y13))#fit is adequate
#run type 3 SS anova
Anova(y13, type="III", test.statistic="LR") #treatment p=0.011 and treatment*minutes p=0.011
#break up data by time point to explore interaction:
#make 30m dataframe
mxlc_GroupMating30 <- mxlc_GroupMating[mxlc_GroupMating$minutes=="30",]
#fit 30m glm
y11 <- glm(spermtransfer~treatment+replicate, family="binomial", data=mxlc_GroupMating30)
summary(y11)
#check model fit using DHARMa
plot(simulateResiduals(fittedModel=y11)) #quantile deviations detected, suggesting fit is inadequate
drop1(y11);summary(y11)#replicate is not significant and removing replicate improves the AIC, so remove replicate
#fit 30m glm without replicate
y11 <- glm(spermtransfer~treatment, family="binomial", data=mxlc_GroupMating30)
summary(y11)
#check model fit using DHARMa
plot(simulateResiduals(fittedModel=y11)) #fit is adequate
#run type 1 SS anova (only one predictor)
anova(y11, test="LRT") #treatment p=0.000414
#make 90m dataframe
mxlc_GroupMating90 <- mxlc_GroupMating[mxlc_GroupMating$minutes=="90",]
#fit 90m glm
y12 <- glm(spermtransfer~treatment+replicate, family="binomial", data=mxlc_GroupMating90)
summary(y12)
#check model fit using DHARMa
plot(simulateResiduals(fittedModel=y12)) #fit is adequate
#run type 3 SS anova
Anova(y12, type="III", test.statistic="LR") #no significant predictors

### Figures
mxlc_GroupMating.prop <- mxlc_GroupMating %>% #get proportions from the binomial data
  group_by(replicate, treatment, minutes) %>%
  summarise(n=n(),
            n.spermtransfer=sum(spermtransfer),
            propmate=n.spermtransfer/n)
mxlc_GroupMating.prop$treatment <- factor(mxlc_GroupMating.prop$treatment, levels=c("mx", "lc"))
mxlc_GroupMating.prop$replicate <- factor(mxlc_GroupMating.prop$replicate, levels=c("one", "two", "three", "four", "five", "six"))
mxlc_GroupMating.prop30 <- mxlc_GroupMating.prop[mxlc_GroupMating.prop$minutes=="30",] #make 30m dataframe
mxlc_GroupMating.prop90 <- mxlc_GroupMating.prop[mxlc_GroupMating.prop$minutes=="90",] #make 90m dataframe
#create fig5a (30min)
fig5a <- boxplot(
  propmate~treatment, data=mxlc_GroupMating.prop30,
  col=c("#FFBF80", "navy"),  # Ensure colors match treatment order
  cex.lab=1.4,
  cex.axis=1.1,
  ylab="Proportion females with sperm stored",
  names=c("MX", "LC"),  # Change x-axis labels
  xlab="30 min",
  ylim=c(0,1.1),
  las=1)
#create fig5b (90min)
fig5b <- boxplot(
  propmate~treatment, data=mxlc_GroupMating.prop90,
  col=c("#FFBF80", "navy"),  # Ensure colors match treatment order
  cex.lab=1.4,
  cex.axis=1.1,
  ylab="Proportion females with sperm stored",
  names=c("MX", "LC"),  # Change x-axis labels
  xlab="30 min",
  ylim=c(0,1.1),
  las=1)
```

##### **Analysis Outputs**

```
y13 <- glm(spermtransfer~treatment*minutes+replicate, family="binomial", data=mxlc_GroupMating)
Anova(y13, type="III", test.statistic="LR")
## Analysis of Deviance Table (Type III tests)
##
## Response: spermtransfer
##                   LR Chisq Df Pr(>Chisq)
## treatment           6.4074  1    0.01136 *
## minutes             1.5729  1    0.20979
## replicate           5.8361  5    0.32250
## treatment:minutes   6.4074  1    0.01136 *
## ---
## Signif. codes:  0 '***' 0.001 '**' 0.01 '*' 0.05 '.' 0.1 ' ' 1
y11 <- glm(spermtransfer~treatment, family="binomial", data=mxlc_GroupMating30)
anova(y11, test="LRT")
## Analysis of Deviance Table
##
## Model: binomial, link: logit
##
## Response: spermtransfer
##
## Terms added sequentially (first to last)
##
##
##           Df Deviance Resid. Df Resid. Dev  Pr(>Chi)
## NULL                         59     79.881
## treatment  1   12.466        58     67.414 0.0004144 ***
## ---
## Signif. codes:  0 '***' 0.001 '**' 0.01 '*' 0.05 '.' 0.1 ' ' 1
y12 <- glm(spermtransfer~treatment+replicate, family="binomial", data=mxlc_GroupMating90)
Anova(y12, type="III", test.statistic="LR")
## Analysis of Deviance Table (Type III tests)
##
## Response: spermtransfer
##           LR Chisq Df Pr(>Chisq)
## treatment    0.000  1     1.0000
## replicate   10.295  5     0.0673 .
## ---
## Signif. codes:  0 '***' 0.001 '**' 0.01 '*' 0.05 '.' 0.1 ' ' 1
# Note about y12 output: LRtreatment=0.00 because sperm transfer success was identical between treatments (21/30 successes for both MX and LC).
```

#### Fig (6A and B): Experiment 1, Competitive mating

##### **R Code**

```
### Load and format data
mxlc_CompMating <- read.table("Fig6_MX_LC_MatingCompetition.csv", sep=",", header=TRUE)
#create table of observed and expected success (success = "winning" the mating competition) for MX and LC
mx.success <- sum(mxlc_CompMating$mx_win==1)
lc.success <- sum(mxlc_CompMating$lc_win==1)
n.trials <- nrow(mxlc_CompMating)
exp <- n.trials/2
tab <- matrix(c(mx.success, exp,
                lc.success, exp),
              nrow=2, byrow=TRUE)
rownames(tab) <- c("MX", "LC")
colnames(tab) <- c("Obs", "Exp")
tab <- as.data.frame(tab) #use for chisq test
#create df of proportions for figures
mxlc_CompMating.prop <- mxlc_CompMating %>% #pivot longer
  pivot_longer(cols=c("mx_win", "lc_win"),
               names_to="treatment",
               values_to="win")
mxlc_CompMating.prop <- mxlc_CompMating.prop %>% #get proportions
  group_by(rep, treatment) %>%
  summarise(n.trials=n(),
            n_win=sum(win),
            proportion=n_win/n.trials)
mxlc_CompMating.prop$rep <- factor(mxlc_CompMating.prop$rep, levels=c("one", "two", "three"))
#group by treatment and get means and SEs
mxlc_CompMating.prop.grouped <- mxlc_CompMating.prop %>%
  group_by(treatment) %>%
  dplyr::summarise(mean_proportion=mean(proportion),
                   se_proportion=sd(proportion) / sqrt(dplyr::n()))  #SE=SD/sqrt(n)
#get cumulative proportions
mxlc_CompMating.prop.grouped <- mxlc_CompMating.prop.grouped %>%
  mutate(ymin=mean_proportion - se_proportion, #get upper error bar
         ymax=mean_proportion + se_proportion) #get lower error bar

### Analyses
#perform chisq test
chisq.test(x=tab$Obs, p=tab$Exp, rescale.p=TRUE) #treatment p=0.4795

### Figures
#create fig6a (stacked bar with replicates combined)
fig6a <- ggplot(
  data=mxlc_CompMating.prop.grouped,
  aes(x="Stacked Bar",
      y=mean_proportion,
      fill=treatment)) +
  geom_bar(stat="identity", width=0.3) +
  geom_errorbar(aes(ymin=ymin[2], ymax=ymax[2]), width=0.1) + #use [2] because MX is on the bottom
  scale_fill_manual(values=c("mx_win"="#FFBF80", "lc_win"="navy"),
                    labels=c("mx_win"="MX", "lc_win"="LC")) +
  labs(x="", y="Proportion winning males", fill="Treatment") +
  ylim(0, 1) +
  theme_minimal() +
  theme(axis.line=element_line(color="black", size=1, linetype=1),
        axis.text.y=element_text(size=14),
        panel.grid.major=element_blank(),
        panel.grid.minor=element_blank(),
        axis.ticks=element_line(size=1),
        axis.text.x=element_blank(),
        axis.ticks.x=element_blank(),
        axis.title.x=element_text(size=20),
        axis.title.y=element_text(size=20),
        legend.title=element_text(size=14),
        legend.text=element_text(size=12))
#create fig6b (stacked bar with replicates separate)
fig6b <- ggplot(
  data=mxlc_CompMating.prop,
  aes(x=rep,
      y=proportion,
      fill=treatment)) +
  geom_bar(stat="identity", position="stack") +
  scale_fill_manual(values=c("mx_win"="#FFBF80", "lc_win"="navy"),
                    labels=c("mx_win"="MX", "lc_win"="LC")) +
  labs(x="Replicate", y="Proportion winning males", fill="Treatment") +
  theme_minimal() +
  theme(axis.line=element_line(color="black", size=1, linetype=1),
        axis.text.x=element_text(size=16),
        axis.text.y=element_text(size=16),
        panel.grid.major=element_blank(),
        panel.grid.minor=element_blank(),
        axis.ticks=element_line(size=1),
        axis.title.x=element_text(size=20),
        axis.title.y=element_text(size=20),
        legend.position="right",
        legend.title=element_text(size=14),
        legend.text=element_text(size=12))
```

##### **Analysis Outputs**

```
chisq.test(x=tab$Obs, p=tab$Exp, rescale.p=TRUE)
##
##  Chi-squared test for given probabilities
##
## data:  tab$Obs
## X-squared = 0.5, df = 1, p-value = 0.4795
```

#### Fig (7, S9, S10): Experiment 1, Bacterial load

##### **R Code**

Note: To assess the effect of treatment and larval presence on bacterial load across time, we ran a mixed measures anova testing effect of days within flask and effects of treatment (MX or LC) and larvae (with larvae or without larvae) between flasks on bacterial load. This model design was chosen because days is the only variable that was measured more than once per flask. For all other factors/variables, each flask was only used for a single level of that factor. Put another way, bacterial treatment was not repeated per flask, each flask had a single independent bacterial treatment applied. This is different than day, because each flask was repeatedly measured (on days 2, 4,6, and 8). We used a mixed measures anova tutorial (<https://www.datanovia.com/en/lessons/mixed-anova-in-r/#three-way-bbw-b>) as a helpful guide for performing this repeated measures analysis.

```
### Load and format data
bl.zero <- read.table("Fig7_MX_LC_BacterialLoad_0day.csv", sep=",", header=TRUE)
bl <- read.table("Fig7_S9_S10_MX_LC_BacterialLoad.csv", sep=",", header=TRUE)
bl$treatment <- factor(bl$treatment, levels=c("mx", "lc", "mx+larvae", "lc+larvae"))
bl$larvae <-as.factor(bl$larvae)
bl$bacttreat <- as.factor(bl$bacttreat)
bl$rep <- as.factor(bl$rep)

### Analyses
#test for outliers
bl %>%
  group_by(bacttreat, larvae, days) %>%
  identify_outliers(logcfuperml) #no extreme outliers
#test for normality:
bl %>%
  group_by(bacttreat, larvae,days) %>%
  shapiro_test(logcfuperml) #only one facet deviates from normality (p=0.028), the rest are adequate
res.aov1 <- anova_test(data=bl, dv=logcfuperml, wid=flask, within=c(days), between=c(larvae, bacttreat))
get_anova_table(res.aov1) #larvae p=1.14e-05; bacttreat p=6.67e-09; days p= 4.00e-03; larvae*bacttreat p=0.020; larvae*days p= 0.044
#create figS9 (interaction plot to visualize 2-way interaction between bacterial treatment and larvae)
figS9 <- interaction.plot(bl$bacttreat, bl$larvae, bl$logcfuperml) #effect is more pronounced in MX vs. LC
#create figS10 (interaction plot to visualize 2-way interaction between larvae and day)
figS10 <- interaction.plot(bl$days, bl$larvae, bl$logcfuperml) #larval presence increases bacterial load until day 8, at which point larval presence decreases bacterial load
#fit anova to test effect of bacterial treatment at each level of larval treatment:
one.way <- bl %>%
  group_by(larvae) %>%
  anova_test(dv=logcfuperml, between=bacttreat) %>%
  get_anova_table() %>%
  adjust_pvalue(method="bonferroni") #larvae absent p.adj=0.00000306; larvae present p.adj=0.000000272
#calculations to understand effect of larvae on change bacterial loads in mx and lc
tapply(bl$cfuperml, bl$treatment, mean) #get the mean cfu of each treatment group
#larvae present
465000000-43741667 # =421258333
421258333/43741667 # =9.630596
43741667+(43741667*9.630596) # =4.65e+08
#larvae absent
271666667-10800000 # =260866667
260866667/10800000 # =24.15432
10800000+(10800000*24.15432) # =271666656

### Figures
#combine bl with day 0 dosing data to make fig7
bl.zero$treatment <- paste0(bl.zero$treatment, ".0")
bl.zero$treatment <- as.factor(bl.zero$treatment)
bl.zero$rep <- as.factor(bl.zero$rep)
bl.combined <- bind_rows(bl, bl.zero)
bl.combined$treatment_x_days <- paste(bl.combined$treatment, bl.combined$days)
#set the ordering in which boxplots appear on x-axis
bl.combined$treatment_x_days <- factor(bl.combined$treatment_x_days,
                                       levels= c("mx.0 0", "lc.0 0",
                                                 "mx 2", "mx 4", "mx 6", "mx 8",
                                                 "lc 2", "lc 4", "lc 6", "lc 8",
                                                 "mx+larvae 2", "mx+larvae 4", "mx+larvae 6", "mx+larvae 8",
                                                 "lc+larvae 2", "lc+larvae 4", "lc+larvae 6", "lc+larvae 8"))
#create fig7
fig7 <- boxplot(
  log10(cfuperml)~treatment_x_days,
  data=bl.combined,
  col=c(rep("grey", 2), rep("#FFE6B3",4), rep("lightblue", 4), rep("#FFBF80",4), rep("navy", 4)),
  ylim=c(3,10),
  las=2,
  ylab="Log10(CFU per ml)",
  xlab="",
  cex.axis=0.7)
```

##### **Analysis Outputs**

```
res.aov1 <- anova_test(data=bl, dv=logcfuperml, wid=flask, within=c(days), between=c(larvae, bacttreat))
get_anova_table(res.aov1)
## ANOVA Table (type II tests)
##
##                  Effect  DFn   DFd       F        p p<.05   ges
## 1                larvae 1.00  8.00  92.380 1.14e-05     * 0.473
## 2             bacttreat 1.00  8.00 632.911 6.67e-09     * 0.860
## 3                  days 1.42 11.37  10.817 4.00e-03     * 0.555
## 4      larvae:bacttreat 1.00  8.00   8.456 2.00e-02     * 0.076
## 5           larvae:days 1.42 11.37   4.629 4.40e-02     * 0.348
## 6        bacttreat:days 1.42 11.37   0.642 4.93e-01       0.069
## 7 larvae:bacttreat:days 1.42 11.37   0.423 5.98e-01       0.046
one.way <- bl %>%
  group_by(larvae) %>%
  anova_test(dv=logcfuperml, between=bacttreat) %>%
  get_anova_table() %>%
  adjust_pvalue(method="bonferroni")
one.way
## # A tibble: 2 × 9
##   larvae  Effect      DFn   DFd     F           p `p<.05`   ges       p.adj
##   <fct>   <chr>     <dbl> <dbl> <dbl>       <dbl> <chr>   <dbl>       <dbl>
## 1 absent  bacttreat     1    22  42.3 0.00000153  *       0.658 0.00000306
## 2 present bacttreat     1    22  57.8 0.000000136 *       0.724 0.000000272
```

### Experiment 2: Comparing development time, adult male longevity, and male wing length between AX, MX and LC individuals.

##### Setup: Load R packages and set plot margins

library(tidyverse)
library(survival)
library(survminer)
library(car)
par(mar=c(10.2,8.2,8.2,4.2))

#### Fig (8A): Experiment 2, Pupation Analysis (AX vs MX vs LC)

##### **R Code**

```
### Load and format data
axmxlc_pup <- read.table("Fig8A_AX_MX_LC_Pupation.csv", sep=",",header=TRUE)
axmxlc_pup$treatment <- factor(axmxlc_pup$treatment, levels=c("ax", "mx", "lc")) #make AX reference
axmxlc_pup$rep <- as.factor(axmxlc_pup$rep)

### Analyses
#fit cox proportional hazards model:
cox5 <- coxph(Surv(daysposthatching, status, type=c('right'))~treatment+rep, data=axmxlc_pup)
cox.zph(cox5, global=T) #coxph global test violated (p=0.045)
#perform logrank test to assess overall effect of treatment on pupation:
survdiff(Surv(daysposthatching,status)~treatment, data=axmxlc_pup) #treatment p=<2e-16
#perform logrank test to assess effect of treatment on pupation within each replicate:
axmxlc_pup_rep1 <- axmxlc_pup %>% dplyr::filter(rep=="one")
axmxlc_pup_rep2 <- axmxlc_pup %>% dplyr::filter(rep=="two")
axmxlc_pup_rep3 <- axmxlc_pup %>% dplyr::filter(rep=="three")
survdiff(Surv(daysposthatching, status)~treatment, data=axmxlc_pup_rep1) #treatment p=<2e-16
survdiff(Surv(daysposthatching, status)~treatment, data=axmxlc_pup_rep2) #treatment p=<2e-16
survdiff(Surv(daysposthatching, status)~treatment, data=axmxlc_pup_rep3) #treatment p=<2e-16
#evaluate pairwise comparisons between treatments
pairwise_survdiff(Surv(daysposthatching,status)~treatment, data=axmxlc_pup, p.adjust.method="BH") #MXvsLC p=0.08; MXvsAX p=<2e-16; AXvsLC p<2e-16

### Figures
#create fig8a
km.axmxlc_pup <- survfit(Surv(daysposthatching,status)~treatment, data=axmxlc_pup) #fit survival object
fig8a <- ggsurvplot(
  km.axmxlc_pup,
  palette=c("#BFD4C9" , "#FFBF80", "navy"),
  conf.int=TRUE,
  font.x= c(16),
  font.y=c(16),
  ylab="Probability of pupation",
  xlab="Days post hatching",
  fun="event",
  risk.table.col="strata",
  legend.labs=c("Axenic", "Monoxenic","Lab Community"))
```

##### **Analysis Outputs**

```
survdiff(Surv(daysposthatching,status)~treatment, data=axmxlc_pup)
## Call:
## survdiff(formula = Surv(daysposthatching, status) ~ treatment,
##     data = axmxlc_pup)
##
##                N Observed Expected (O-E)^2/E (O-E)^2/V
## treatment=ax 108      104    179.2      31.5     260.8
## treatment=mx  72       72     32.6      47.7     100.0
## treatment=lc  72       72     36.3      35.2      73.8
##
##  Chisq= 263  on 2 degrees of freedom, p= <2e-16
survdiff(Surv(daysposthatching, status)~treatment, data=axmxlc_pup_rep1) # replicate 1
## Call:
## survdiff(formula = Surv(daysposthatching, status) ~ treatment,
##     data = axmxlc_pup_rep1)
##
##               N Observed Expected (O-E)^2/E (O-E)^2/V
## treatment=ax 36       35    60.31     10.62      84.3
## treatment=mx 24       24     9.92     19.97      39.2
## treatment=lc 24       24    12.76      9.89      20.0
##
##  Chisq= 87.6  on 2 degrees of freedom, p= <2e-16
survdiff(Surv(daysposthatching, status)~treatment, data=axmxlc_pup_rep2) # replicate 2
## Call:
## survdiff(formula = Surv(daysposthatching, status) ~ treatment,
##     data = axmxlc_pup_rep2)
##
##               N Observed Expected (O-E)^2/E (O-E)^2/V
## treatment=ax 36       35     60.0      10.4      83.2
## treatment=mx 24       24     11.1      15.0      30.9
## treatment=lc 24       24     12.0      12.1      24.6
##
##  Chisq= 83.7  on 2 degrees of freedom, p= <2e-16
survdiff(Surv(daysposthatching, status)~treatment, data=axmxlc_pup_rep3) # replicate 3
## Call:
## survdiff(formula = Surv(daysposthatching, status) ~ treatment,
##     data = axmxlc_pup_rep3)
##
##               N Observed Expected (O-E)^2/E (O-E)^2/V
## treatment=ax 36       34     58.5      10.3      90.5
## treatment=mx 24       24     11.5      13.4      30.4
## treatment=lc 24       24     12.0      12.1      27.9
##
##  Chisq= 90.5  on 2 degrees of freedom, p= <2e-16
pairwise_survdiff(Surv(daysposthatching,status)~treatment, data=axmxlc_pup, p.adjust.method="BH")
##
##  Pairwise comparisons using Log-Rank test
##
## data:  axmxlc_pup and treatment
##
##    ax     mx
## mx <2e-16 -
## lc <2e-16 0.088
##
## P value adjustment method: BH
```

#### Fig (8B): Experiment 2, Eclosion Analysis (AX vs MX vs LC)

##### **R Code**

```
### Load and format data
axmxlc_eclo <- read.table("Fig8B_AX_MX_LC_Eclosion.csv", sep=",", header=TRUE)
axmxlc_eclo$treatment <- factor(axmxlc_eclo$treatment, levels=c("ax", "mx", "lc")) #set reference level and order
axmxlc_eclo$rep <- as.factor(axmxlc_eclo$rep)

### Analyses
#fit cox proportional hazards model:
cox6 <- coxph(Surv(daysposthatching, status, type=c('right'))~treatment+rep, data=axmxlc_eclo)
cox.zph(cox6, global=T) #coxph assumptions met
summary(cox6)
anova(cox6) #treatment p=<2e-16
#evaluate pairwise comparisons between treatments
pairwise_survdiff(Surv(daysposthatching,status)~treatment, data=axmxlc_eclo, p.adjust.method="BH") #MXvsLC p=0.2; MXvsAX p=<2e-16; AXvsLC p<2e-16

### Figures
#create fig8b
km.axmxlc_eclo <- survfit(Surv(daysposthatching,status)~treatment, data=axmxlc_eclo) #fit survival object
fig8b <- ggsurvplot(
  km.axmxlc_eclo,
  palette=c("#BFD4C9" , "#FFBF80", "navy"),
  conf.int=TRUE,
  font.x= c(16),
  font.y=c(16),
  ylab="Probability of eclosion",
  xlab="Days post hatching",
  fun="event",
  risk.table.col="strata",
  legend.labs=c("Axenic", "Monoxenic","Lab Community"))
```

##### **Analysis Outputs**

```
cox6 <- coxph(Surv(daysposthatching, status, type=c('right'))~treatment+rep, data=axmxlc_eclo)
anova(cox6)
## Analysis of Deviance Table
##  Cox model: response is Surv(daysposthatching, status, type = c("right"))
## Terms added sequentially (first to last)
##
##             loglik    Chisq Df Pr(>|Chi|)
## NULL      -1138.52
## treatment  -968.89 339.2667  2     <2e-16 ***
## rep        -968.68   0.4107  2     0.8144
## ---
## Signif. codes:  0 '***' 0.001 '**' 0.01 '*' 0.05 '.' 0.1 ' ' 1
pairwise_survdiff(Surv(daysposthatching,status)~treatment, data=axmxlc_eclo, p.adjust.method="BH")
##
##  Pairwise comparisons using Log-Rank test
##
## data:  axmxlc_eclo and treatment
##
##    ax     mx
## mx <2e-16 -
## lc <2e-16 0.2
##
## P value adjustment method: BH
```

#### Fig (8C): Experiment 2, Survival Analysis in Sugar-Fed (AX vs MX vs LC)

##### **R Code**

```
### Load and format data
axmxlc_death <- read.table("Fig8C_AX_MX_LC_Survival.csv", sep=",", header=TRUE)
axmxlc_death$treatment <- factor(axmxlc_death$treatment, levels=c("ax", "mx", "lc")) #make AX reference
axmxlc_death$rep <- as.factor(axmxlc_death$rep)

### Analyses
#fit cox proportional hazards model:
cox7 <- coxph(Surv(daysposteclosion, status, type=c('right'))~treatment+rep, data=axmxlc_death)
cox.zph(cox7, global=T) #coxph assumption violated for treatment
#perform logrank test to assess overall effect of treatment on survival:
survdiff(Surv(daysposteclosion, status)~treatment, data=axmxlc_death) #treatment p=2e-05
#perform logrank test to assess effect of treatment on eclosion within each replicate:
axmxlc_death_rep4 <- axmxlc_death %>% dplyr::filter(rep=="four")
axmxlc_death_rep5 <- axmxlc_death %>% dplyr::filter(rep=="five")
survdiff(Surv(daysposteclosion, status)~treatment, data=axmxlc_death_rep4) #treatment p=3e-04
survdiff(Surv(daysposteclosion, status)~treatment, data=axmxlc_death_rep5) #treatment p=0.009
#evaluate pairwise comparisons between treatments
pairwise_survdiff(Surv(daysposteclosion,status)~treatment, data=axmxlc_death, p.adjust.method="BH") #MXvsLC p=0.041; MXvsAX p=0.0078; AXvsLC p=1.3e-05

### Figures
#create fig8c
km.axmxlc_death <- survfit(Surv(daysposteclosion,status)~treatment, data=axmxlc_death) #fit survival object
fig8c <- ggsurvplot(
  km.axmxlc_death,
  conf.int=TRUE,
  ylab="Survival probability",
  xlab="Days post eclosion",
  font.x=c(16),
  font.y=c(16),
  palette=c("#BFD4C9","#FFBF80","navy"),
  legend.labs=
    c("Axenic", "Monoxenic", "Lab Community"))
```

##### **Analysis Outputs**

```
survdiff(Surv(daysposteclosion, status)~treatment, data=axmxlc_death)
## Call:
## survdiff(formula = Surv(daysposteclosion, status) ~ treatment,
##     data = axmxlc_death)
##
##               N Observed Expected (O-E)^2/E (O-E)^2/V
## treatment=ax 22       22     37.0     6.065    15.009
## treatment=mx 25       25     22.2     0.353     0.573
## treatment=lc 25       25     12.8    11.559    16.293
##
##  Chisq= 21.7  on 2 degrees of freedom, p= 2e-05
survdiff(Surv(daysposteclosion, status)~treatment, data=axmxlc_death_rep4)
## Call:
## survdiff(formula = Surv(daysposteclosion, status) ~ treatment,
##     data = axmxlc_death_rep4)
##
##               N Observed Expected (O-E)^2/E (O-E)^2/V
## treatment=ax 11       11    19.73      3.86     15.60
## treatment=mx 10       10     6.39      2.04      3.24
## treatment=lc 10       10     4.88      5.38      7.96
##
##  Chisq= 16.4  on 2 degrees of freedom, p= 3e-04
survdiff(Surv(daysposteclosion, status)~treatment, data=axmxlc_death_rep5)
## Call:
## survdiff(formula = Surv(daysposteclosion, status) ~ treatment,
##     data = axmxlc_death_rep5)
##
##               N Observed Expected (O-E)^2/E (O-E)^2/V
## treatment=ax 11       11    17.29    2.2892    4.4579
## treatment=mx 15       15    15.63    0.0251    0.0441
## treatment=lc 15       15     8.08    5.9201    8.3127
##
##  Chisq= 9.4  on 2 degrees of freedom, p= 0.009
pairwise_survdiff(Surv(daysposteclosion,status)~treatment, data=axmxlc_death, p.adjust.method="BH")
##
##  Pairwise comparisons using Log-Rank test
##
## data:  axmxlc_death and treatment
##
##    ax      mx
## mx 0.0078  -
## lc 1.3e-05 0.0412
##
## P value adjustment method: BH
```

#### Fig (8D): Experiment 2, Wing length (AX vs MX vs LC)

##### **R Code**

```
### Load and format data
axmxlc_maleWL <- read.table("Fig8D_AX_MX_LC_WingLength.csv", sep=",", header=TRUE)
axmxlc_maleWL$treatment <- factor(axmxlc_maleWL$treatment, levels=c("ax", "mx", "lc")) #set reference level and order
axmxlc_maleWL$replicate <- as.factor(axmxlc_maleWL$replicate)

### Analyses
#assess effect of treatment and replicate on winglength with all replicates combined:
#set contrasts for type III sums of squares
contrasts(axmxlc_maleWL$treatment) <- contr.sum
contrasts(axmxlc_maleWL$replicate) <- contr.sum
y3 <- lm(malewinglength~treatment+replicate, data=axmxlc_maleWL)
#check model fit
plot(y3) #fit is adequate
summary(y3)
#run type 3 SS anova
Anova(y3, type="III", test.statistic="F") #treatment p=<2e-16
#Tukey’s HSD test to evaluate pairwise comparisons between treatment groups:
TukeyHSD(aov(y3), "treatment") #MXvsLC p.adj=0.249; MXvsAX p.adj=0.00 (appx); AXvsLC p.adj=0.00 (appx)

### Figures
#create fig8d
fig8d <- boxplot(
  malewinglength~treatment, data=axmxlc_maleWL,
  col=c("#F2F2F2","#FFBF80", "navy"),
  cex.lab=1.5,
  cex.axis=1.1,
  ylab="Male wing length (mm)",
  names=c("AX", "MX", "LC"),  # Change x-axis labels
  xlab="",
  las=1)  # Rotate y-axis numbers to vertical
```

##### **Analysis Outputs**

```
y3 <- lm(malewinglength~treatment+replicate, data=axmxlc_maleWL)
Anova(y3, type="III", test.statistic="F")
## Anova Table (Type III tests)
##
## Response: malewinglength
##              Sum Sq Df    F value Pr(>F)
## (Intercept) 213.800  1 35891.5771 <2e-16 ***
## treatment     0.982  2    82.4144 <2e-16 ***
## replicate     0.010  1     1.6828 0.2004
## Residuals     0.304 51
## ---
## Signif. codes:  0 '***' 0.001 '**' 0.01 '*' 0.05 '.' 0.1 ' ' 1
TukeyHSD(aov(y3), "treatment")
##   Tukey multiple comparisons of means
##     95% family-wise confidence level
##
## Fit: aov(formula = y3)
##
## $treatment
##             diff         lwr       upr    p adj
## mx-ax 0.25802632  0.19757863 0.3184740 0.000000
## lc-ax 0.29961300  0.23741284 0.3618132 0.000000
## lc-mx 0.04158669 -0.02061347 0.1037868 0.249061
```

### Experiment 3: Absence of microbiota in adult stage

##### Setup: Load R packages and set plot margins

library(survival)
library(survminer)
par(mar=c(10.2,8.2,8.2,4.2))

#### Fig (9): Experiment 3, Survival Analysis, ampicillin treatment

##### **R Code**

```
### Load and format data
ampEcoli_death <- read.table("Fig9_MX_Ampicillin_Survival.csv", sep=",", header=TRUE)
ampEcoli_death$treatment <- as.factor(ampEcoli_death$treatment)
ampEcoli_death$rep <- as.factor(ampEcoli_death$rep)
ampEcoli_death$amp <- as.factor(ampEcoli_death$amp)
ampEcoli_death$e.coli <- as.factor(ampEcoli_death$e.coli)

### Analyses
#fit cox proportional hazards model:
cox8 <- coxph(Surv(daypostinfection, status, type=c('right'))~treatment+rep, data=ampEcoli_death)
cox.zph(cox8, global=T) #coxph assumptions met
summary(cox8)
anova(cox8) #treatment p=0.0008909

### Figures
#create fig9
km.ampEcoli_death <- survfit(Surv(daypostinfection,status)~treatment, data=ampEcoli_death) #fit survival object
fig9 <- ggsurvplot(
  km.ampEcoli_death,
  conf.int=TRUE,
  ylab="Survival probability",
  xlab= expression("Days post " * italic("E. coli") * " re-introduction"),
  font.x=c(20),
  font.y=c(20),
  palette=c("green","darkgreen", "#FFBF80", "tan4"),
  legend.labs=
    expression("AX_Amp", "AX_Amp+E. coli-fed", "MX", "MX_E. coli-fed"),
  font.legend=c(13),
  font.tickslab=c(16))  # Adjust x and y axis tick label size
```

##### **Analysis Outputs**

```
cox8 <- coxph(Surv(daypostinfection, status, type=c('right'))~treatment+rep, data=ampEcoli_death)
anova(cox8)
## Analysis of Deviance Table
##  Cox model: response is Surv(daypostinfection, status, type = c("right"))
## Terms added sequentially (first to last)
##
##            loglik   Chisq Df Pr(>|Chi|)
## NULL      -300.30
## treatment -300.12  0.3647  3  0.9474389
## rep       -294.60 11.0417  1  0.0008909 ***
## ---
## Signif. codes:  0 '***' 0.001 '**' 0.01 '*' 0.05 '.' 0.1 ' ' 1
pairwise_survdiff(Surv(daypostinfection,status)~treatment, data=ampEcoli_death, p.adjust.method="none")
##
##  Pairwise comparisons using Log-Rank test
##
## data:  ampEcoli_death and treatment
##
##             amp200 amp200ecoli ampneg
## amp200ecoli 0.85   -           -
## ampneg      0.53   0.41        -
## ampnegecoli 0.97   0.97        0.79
##
## P value adjustment method: none
```

#### Fig (S11): Experiment 3, Pupation Analysis, ampicillin treatment

##### **R Code**

```
### Load and format data
amp_pup <- read.table("FigS11_MX_Ampicillin_Pupation.csv", sep=",", header=TRUE)
amp_pup$treatment<- as.factor(amp_pup$treatment) #treatment is ampicillin treatment
amp_pup$rep <- as.factor(amp_pup$rep)

### Analyses
#fit cox proportional hazards model:
cox9 <- coxph(Surv(dayposthatching, status, type=c('right'))~treatment+rep, data=amp_pup)
cox.zph(cox9, global=T) #coxph assumptions violated for rep
#fit cox proportional hazards model without rep:
cox9 <- coxph(Surv(dayposthatching, status, type=c('right'))~treatment, data=amp_pup)
cox.zph(cox9, global=T) #coxph assumptions met
summary(cox9)
anova(cox9) #treatment p=0.5949

### Figures
#create figS11
km.amp_pup <- survfit(Surv(dayposthatching,status)~treatment, data=amp_pup) #fit survival object
figS11 <- ggsurvplot(
  km.amp_pup,
  fun="event",
  conf.int=TRUE,
  ylab="Probability of pupation",
  xlab="Days post hatching",
  break.time.by=2,
  font.x=c(20),
  font.y=c(20),
  palette=c("darkgreen","#FFBF80"),
  legend.labs=
    c("Ampicillin treated", "Untreated"),
  font.legend=c(13),
  font.tickslab=c(16))
```

##### **Analysis Outputs**

```
cox9 <- coxph(Surv(dayposthatching, status, type=c('right'))~treatment, data=amp_pup)
anova(cox9)
## Analysis of Deviance Table
##  Cox model: response is Surv(dayposthatching, status, type = c("right"))
## Terms added sequentially (first to last)
##
##            loglik  Chisq Df Pr(>|Chi|)
## NULL      -2039.3
## treatment -2039.2 0.2828  1     0.5949
```

#### Fig (S12): Experiment 3, Eclosion Analysis, ampicillin treatment

##### **R Code**

```
### Load and format data
amp_eclo <- read.table("FigS12_MX_Ampicillin_Eclosion.csv", sep=",", header=TRUE)
amp_eclo$treatment<- as.factor(amp_eclo$treatment) #treatment is ampicillin treatment
amp_eclo$rep <- as.factor(amp_eclo$rep)

### Analyses
#fit cox proportional hazards model:
cox10 <- coxph(Surv(dayposthatching, status, type=c('right'))~treatment+rep, data=amp_eclo)
cox.zph(cox10, global=T) #coxph assumptions violated for rep
#fit cox proportional hazards model without rep:
cox10 <- coxph(Surv(dayposthatching, status, type=c('right'))~treatment, data=amp_eclo)
cox.zph(cox10, global=T) #coxph assumptions met
summary(cox10)
anova(cox10) #treatment p=0.5823

### Figures
#create figS12
km.amp_eclo <- survfit(Surv(dayposthatching,status)~treatment, data=amp_eclo) #fit survival object
figS12 <- ggsurvplot(
  km.amp_eclo,
  fun="event",
  conf.int=TRUE,
  ylab="Probability of eclosion",
  xlab="Days post hatching",
  break.time.by=2,
  font.x=c(20),
  font.y=c(20),
  palette=c("darkgreen","#FFBF80"),
  legend.labs=
    c("Ampicillin treated", "Untreated"),
  font.legend=c(13),
  font.tickslab=c(16))
```

##### **Analysis Outputs**

```
cox10 <- coxph(Surv(dayposthatching, status, type=c('right'))~treatment, data=amp_eclo)
anova(cox10)
## Analysis of Deviance Table
##  Cox model: response is Surv(dayposthatching, status, type = c("right"))
## Terms added sequentially (first to last)
##
##            loglik  Chisq Df Pr(>|Chi|)
## NULL      -1944.5
## treatment -1944.3 0.3025  1     0.5823
```
