## Supplementary figures S1-S12 for "The microbiota impacts life history traits and mating success in male *Aedes aegypti* mosquitoes"

**Supplementary figures and tables**

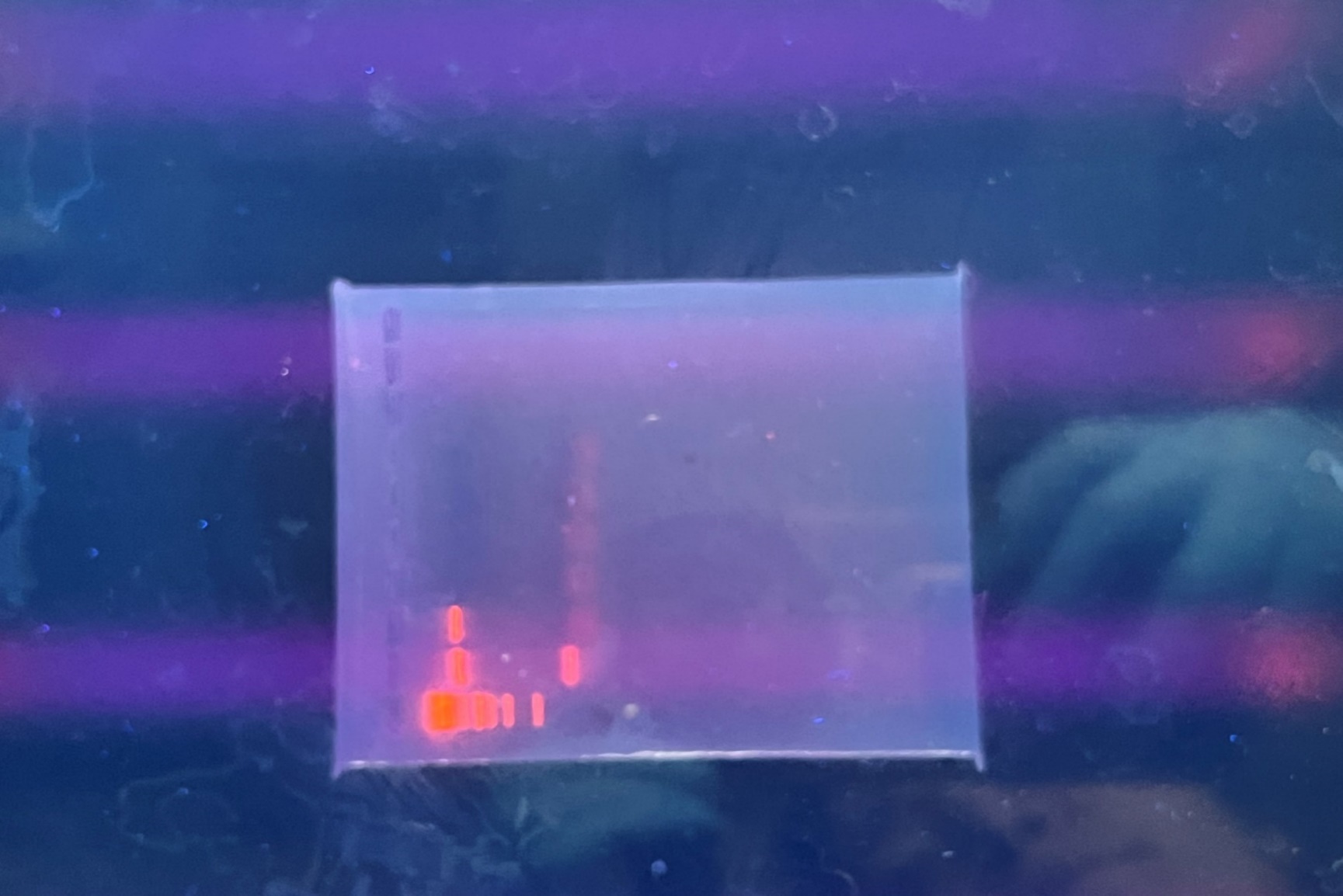

**1	2	3	4	5	6	7**

**Target size**

**~1.5 kb**

1- 1 kb DNA Ladder (Life Technologies, USA)
2- Positive control (*E. coli*)
3- Conventional adult
4- Axenic larvae
5- Axenic adult
6- Extraction control (extraction kit eluate used as template)
7- PCR control (no template control)

**Fig. S1.** **Initial verification of sterilization procedure and maintenance of axenic larvae and adult males via PCR.** A gel electrophoresis image showing the sterility of axenic larvae and axenic adult males reared in a bacteria-free environment, compared to conventional adults (CN) isolated from our laboratory colony. Amplification of the bacterial 16S rRNA gene was performed using the universal primers fD1/fD2 and rP1/rP2 (Weisburg et al., 1991), resulting in an expected amplicon size of approximately 1.5 kb. The DNA extraction was performed using the DNeasy Blood & Tissue kit (Qiagen, USA) according to the manufacturer’s instructions. PCR reagents were prepared as follows: 0.5 μL of each primer at a concentration of 10 μM was added to 12.5 μL of MyTaq Red master mix (Fisher Scientific, USA), and then molecular grade water was added to a final volume of 25 μL. The PCR cycling conditions were as follows: 95°C for 3 minutes, followed by 30 cycles of (95°C for 30 seconds, 55°C for 30 seconds, 72°C for 30 seconds), then 72°C for 10 minutes, and finally a 10°C hold.

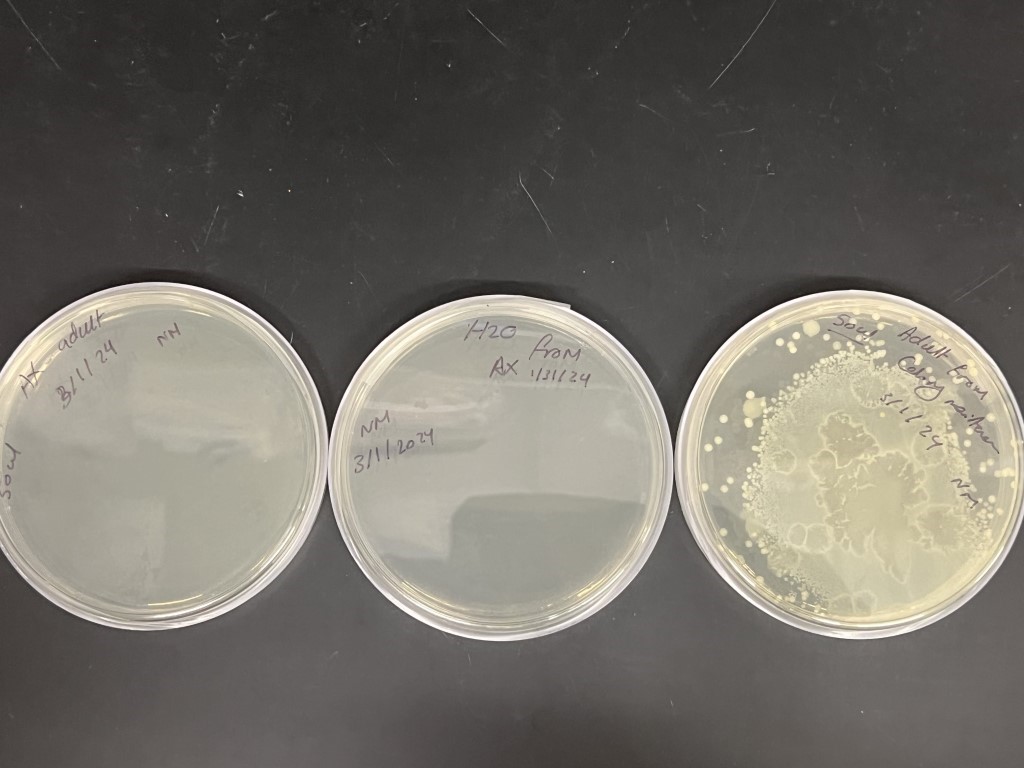

**CN Adult**

**AX larval water**

**AX adult**

**Fig. S2.** **Initial verification of sterilization procedure and maintenance of axenic larvae and adult males via culturing on LB agar medium.** Representative LB agar plates showing the sterility of axenic (AX) adult males reared in a bacteria-free environment and axenic larval water, compared to conventional adults (CN) isolated from our laboratory colony. The culturing was carried out by inoculating LB plates with 50 µL of axenic larval water or adult male homogenate. To make the male homogenate, one adult AX or CN male was homogenized in 150 µL of 1X PBS. Three males were homogenzied and plated per sterility check. The plates were incubated at room temperature for one week to assess sterility.

**A**

**B**

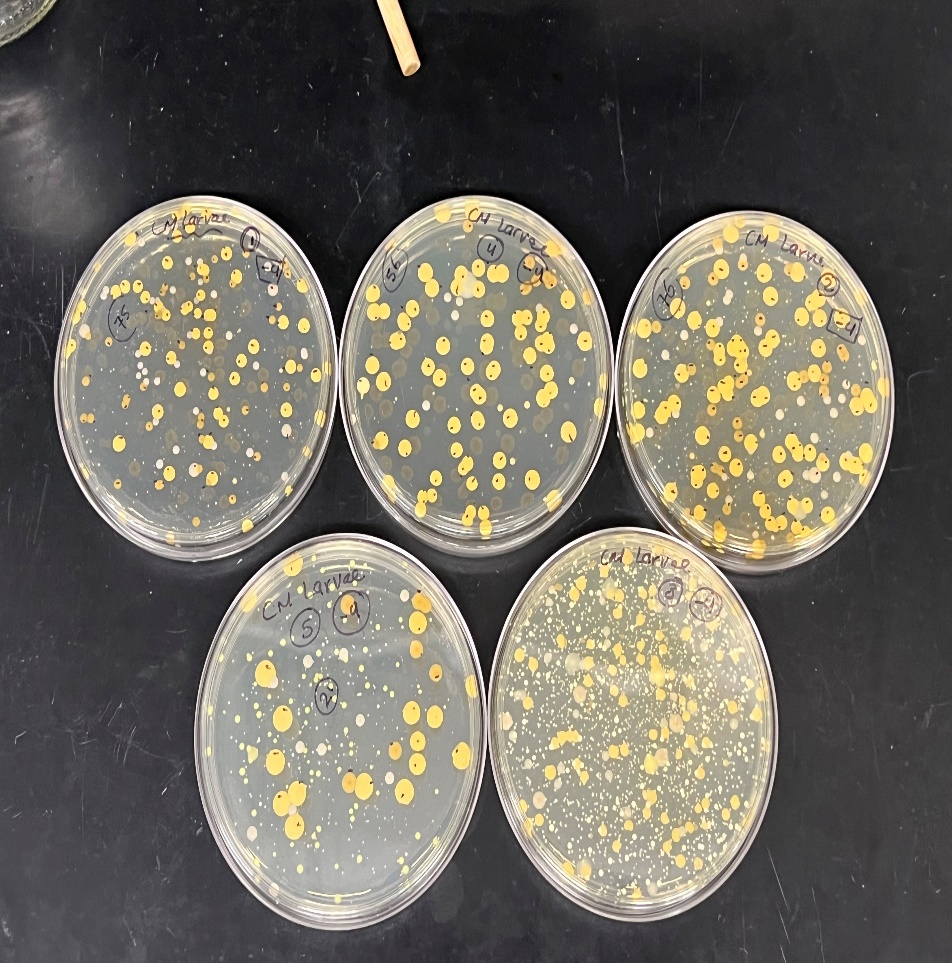

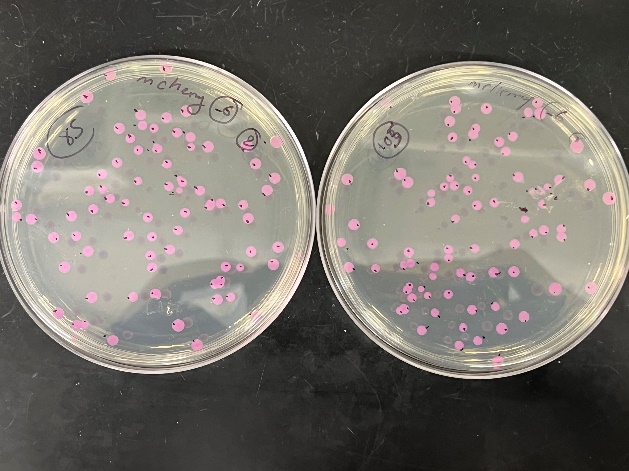

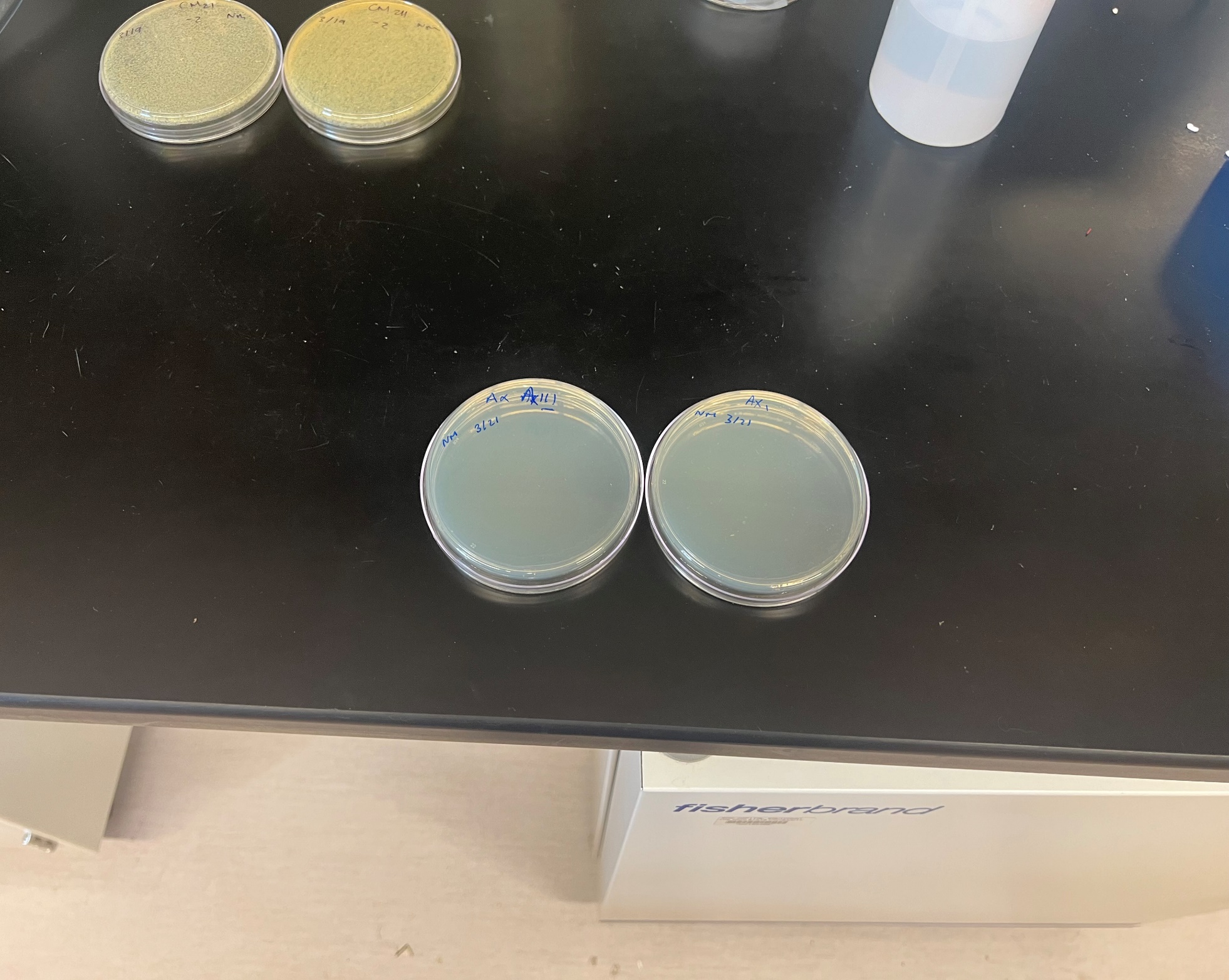

**C**

**Fig. S3.** **Verification of the success of microbiota treatment for each experiment via culturing homogenized adult males on LB agar medium.** Representative images of males homogenized and cultured on LB agar plates. This was performed for each experiment to assess the success of each microbiota treatment. Three adult males from each treatment were individually homogenzied in separate tubes containing 150 µL of 1X PBS. 100 µL of each homogenate was spread on LB agar and incubated at room temperature for one week. (A) Axenic treatment (AX) showed no bacterial growth, (B) monoxenic treatment (MX) showed a single bacterial species, *E. coli* S17 pPROBE-mCherry, and (C)
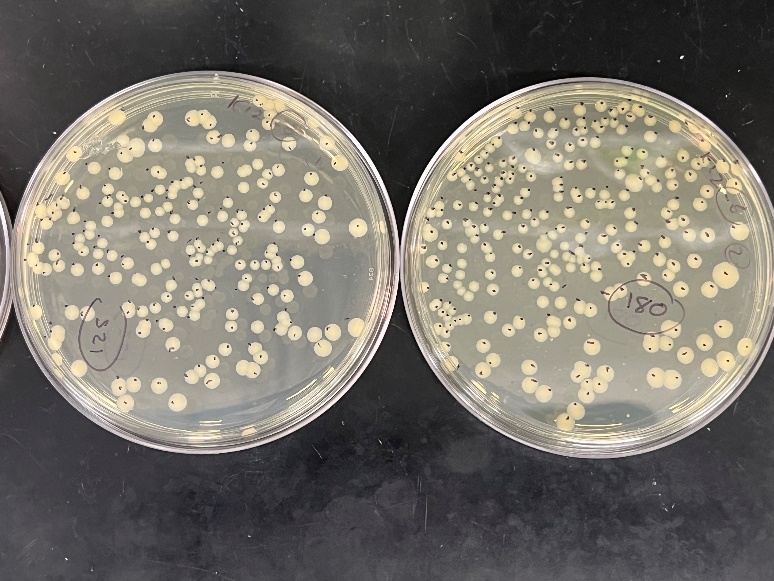
lab community treatment (LC) showed a mixed microbial community.

A

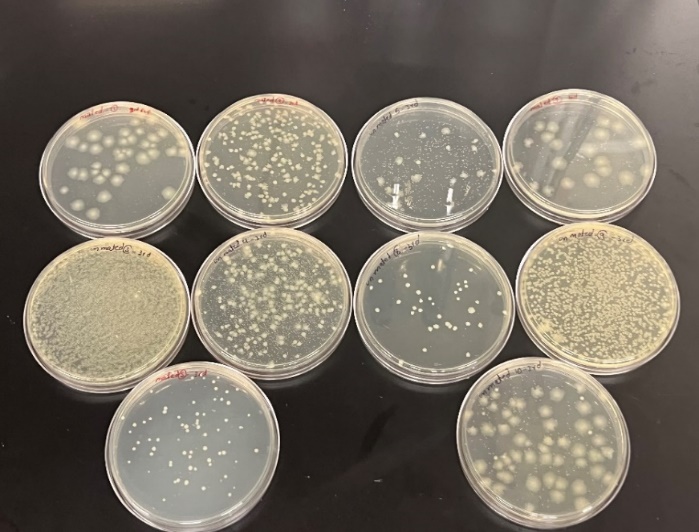

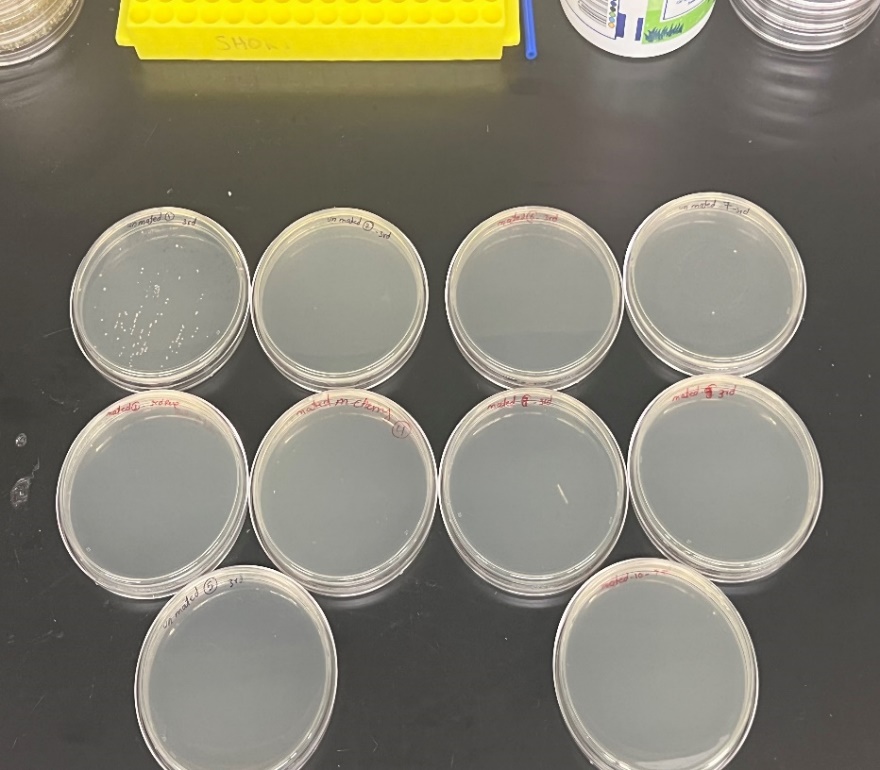

B

**Fig. S4.** **Identification of microbiota treatment (MX or LC) of males upon completion of competitive mating trials.** Representative images of the homogenates of males from competitive mating experiments cultured on LB agar plates. (A) Males that yielded zero or very few colonies were determined to be from the MX group. (B) Males that yielded relatively high numbers of colonies composed of multiple different colony types were determined to be from the LC group.

A

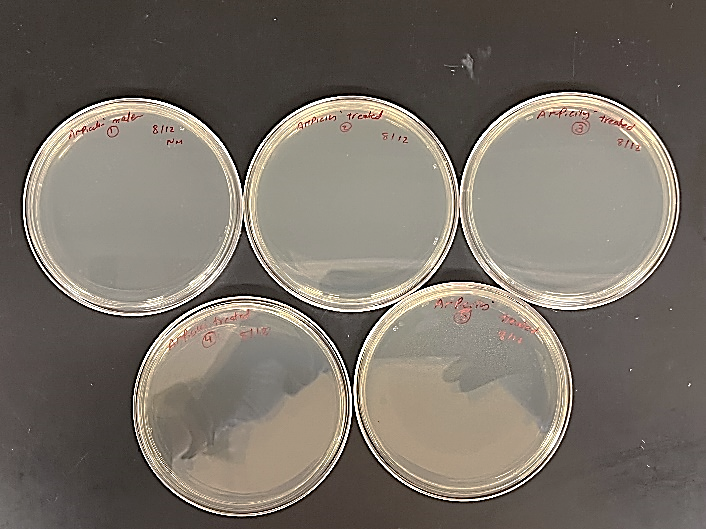

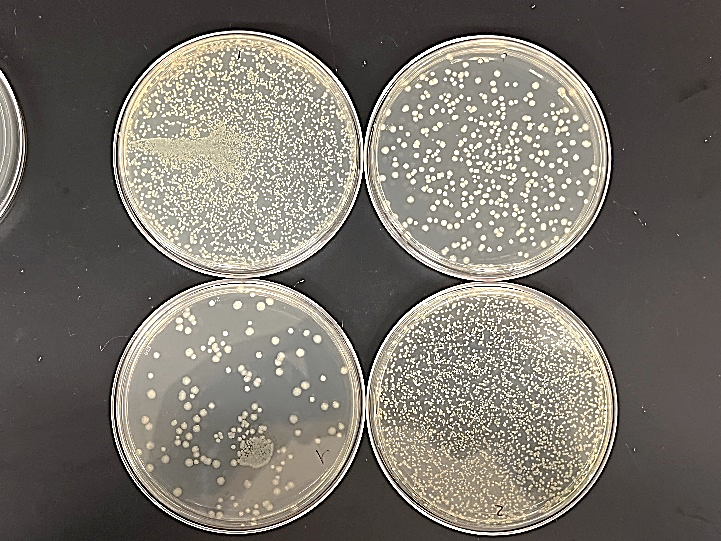

B

**Fig. S5.** **Verification of successful ampicillin treatment via culturing homogenized adult males on LB agar medium.** Representative images of males homogenized and cultured on LB agar plates. This was performed for each experiment to assess the success of ampicillin treatment. Inividual adult males were homogenized in 150 µL 1X PBS and 100 µL of homogenate was cultured on LB agar at room tempature for one week. (A) Ampicillin treated males (AX_amp_) showed no bacterial growth, (B) Non-ampicillin treated males (MX) showed a single bacterial species, *E. coli* K12.

**Path analysis:**

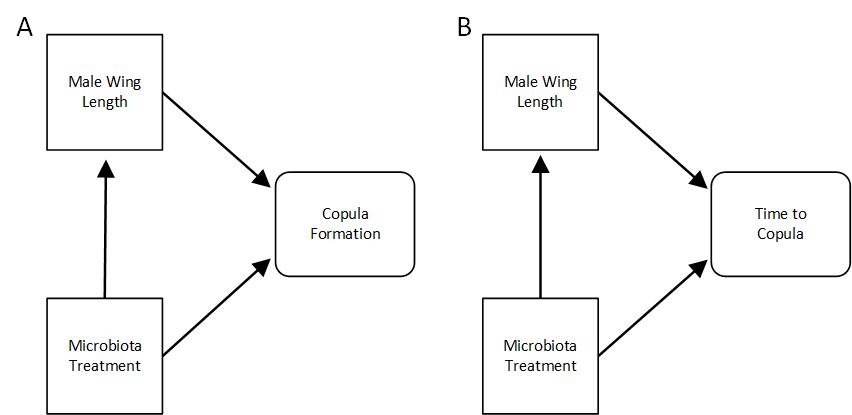

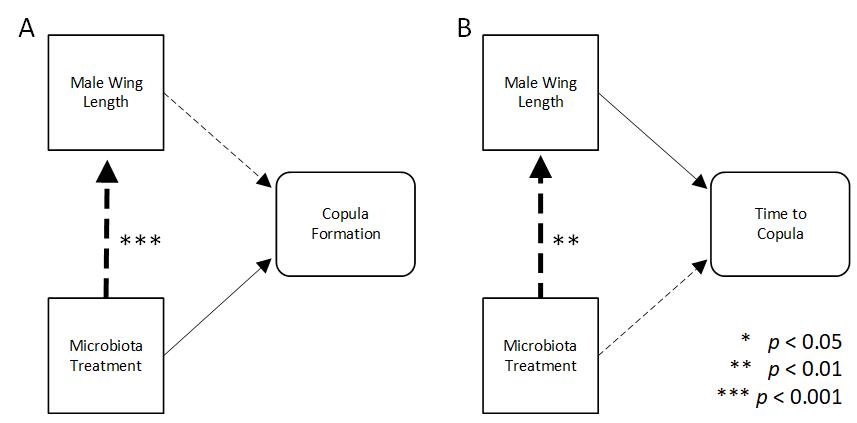

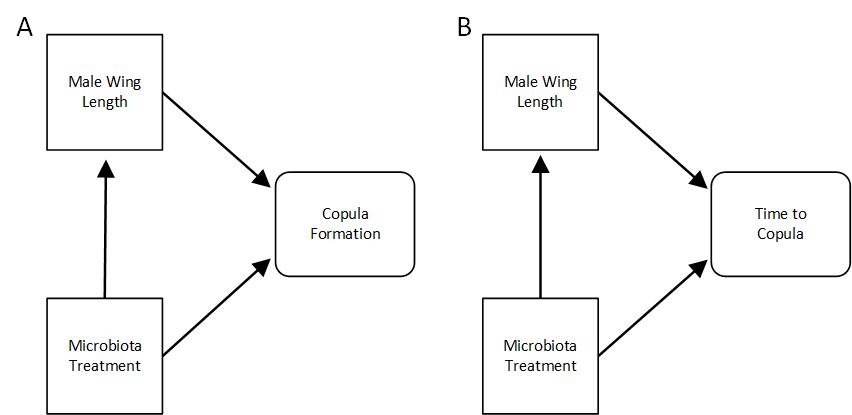

**Fig. S6. Path analysis for time to copula formation.** We conducted a path analysis to determine whether there is an indirect relationship between microbiota treatment and time to copula formation and whether this relationship is mediated by male wing length according to the conceptual model (A). We found that direct relationships were not detected among time to copula formation and either male wing length or microbiota treatment (Table S1). Male wing length was negatively affected by microbiota treatment (B; Std. Est. = -0.38, p = 0.002), but this relationship did not indirectly impact time to copula formation (Table S1). The CFI and RMSEA values were 1.00 and 0.00, respectively, indicating the model is a good approximate fit to the data, but the Chi-square test indicated a poor fit (Χ2 test, p = 0.005), likely due to sample size.

**Table S1. Path analysis model coefficients for the response variable time to copula formation.**

| Pathway | Response  variable | Predictor  variable | Estimate | Std  error | *z*-value | *p*-value | Std  estimate |
| --- | --- | --- | --- | --- | --- | --- | --- |
| Direct |  |  |  |  |  |  |  |
|  | Time to copula ~ | Male wing length | 1.87 | 1.23 | 1.52 | 0.128 | 0.25 |
|  |  | Microbiota treatment | -0.10 | 0.29 | -0.36 | 0.717 | -0.05 |
|  | Male wing length ~ | Microbiota treatment | -0.10 | 0.03 | -3.07 | 0.002 | -0.38 |
| Indirect |  |  |  |  |  |  |  |
| Microbiota treatment 🡪 male wing length 🡪 time to copula | | | -0.19 | 0.12 | -1.52 | 0.127 | -0.09 |

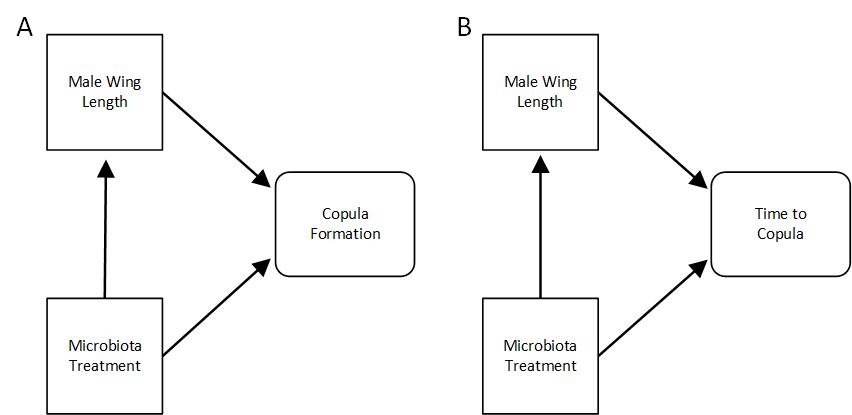

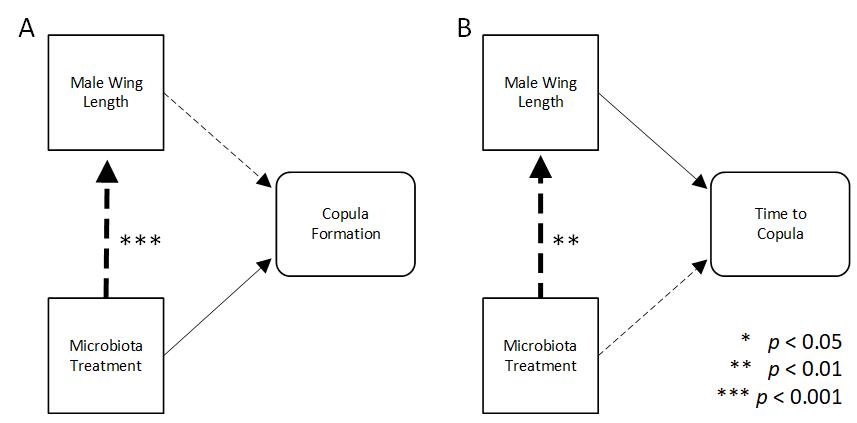

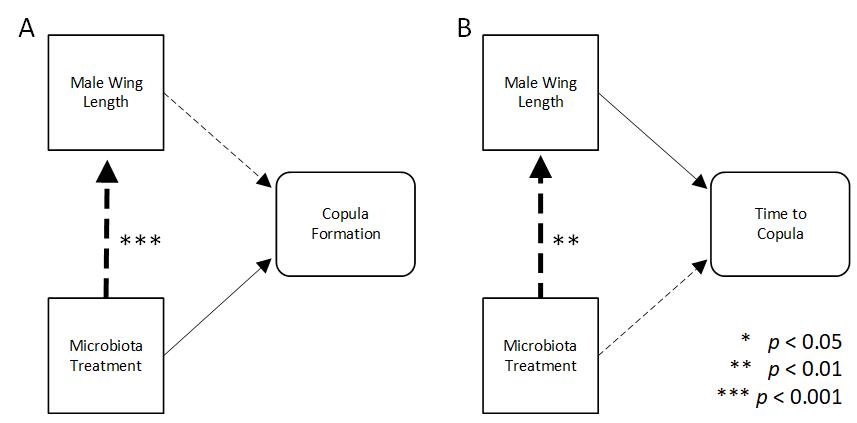

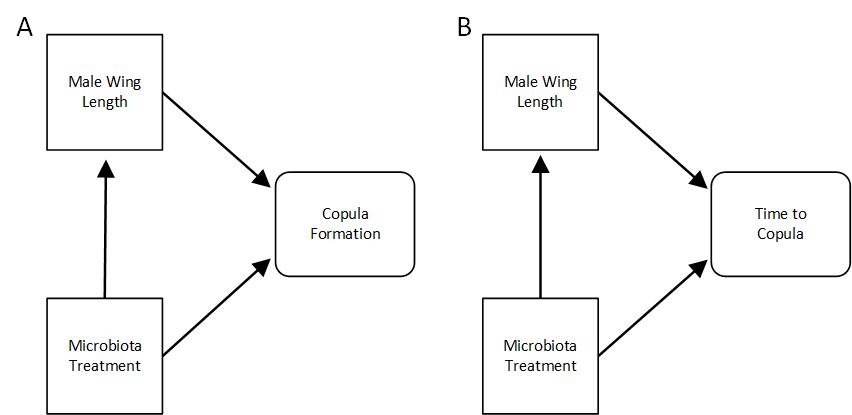

**Fig. S7. Path analysis for copulation success.** We conducted a path analysis to determine whether there is an indirect relationship between microbiota treatment and copulation success and whether this relationship is mediated by male wing length according to the conceptual model (A). We found that copulation success was unaffected by male wing length and microbiota treatment (Table S2). Male wing length was negatively affected by microbiota treatment (B; Std. Est. = -0.39, p < 0.001), but this relationship did not indirectly impact copulation success (Table S2). The path analysis model was consistent with the data (Χ2 test, p = 0.772), and CFI and RMSEA values were 1.00 and 0.00, respectively, indicating good model fit.

**Table S2. Path analysis model coefficients for the response variable copulation success.**

| Pathway | Response  variable | Predictor  variable | Estimate | Std  error | *z*-value | *p*-value | Std  estimate |
| --- | --- | --- | --- | --- | --- | --- | --- |
| Direct |  |  |  |  |  |  |  |
|  | Copula formation ~ | Male wing length | -0.39 | 1.28 | -0.31 | 0.756 | -0.04 |
|  |  | Microbiota treatment | 0.12 | 0.33 | 0.37 | 0.706 | 0.06 |
|  | Male wing length ~ | Microbiota treatment | -0.09 | 0.02 | -3.66 | 0.000 | -0.39 |
| Indirect |  |  |  |  |  |  |  |
| Microbiota treatment 🡪 male wing length 🡪 copula formation | | | 0.03 | 0.12 | 0.29 | 0.765 | 0.01 |

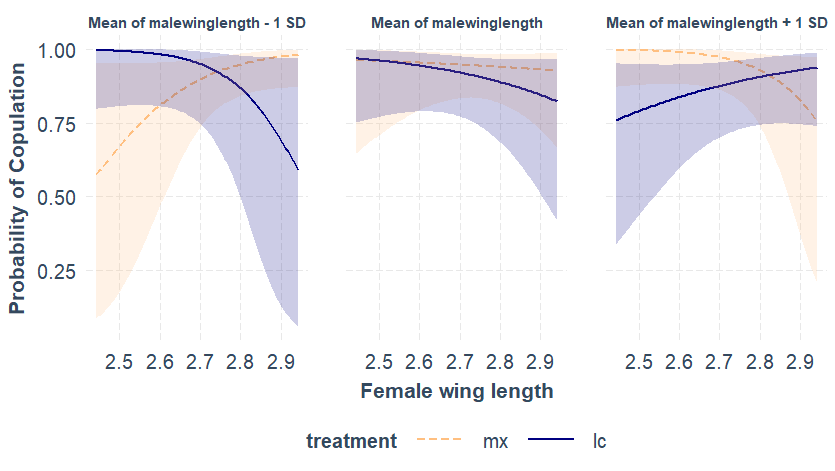

**Fig. S8. Visualization of the 3-way interaction between microbiota treatment, male wing length, and female wing length in predicting copulation success.** The 3-way interaction plot generated using interact_plot() from the R package *interactions* shows that small MX males were more likely to copulate when females were large, but small LC males were more likely to copulate when females were small (left-most panel) and this was the opposite for large males (right-most panel).

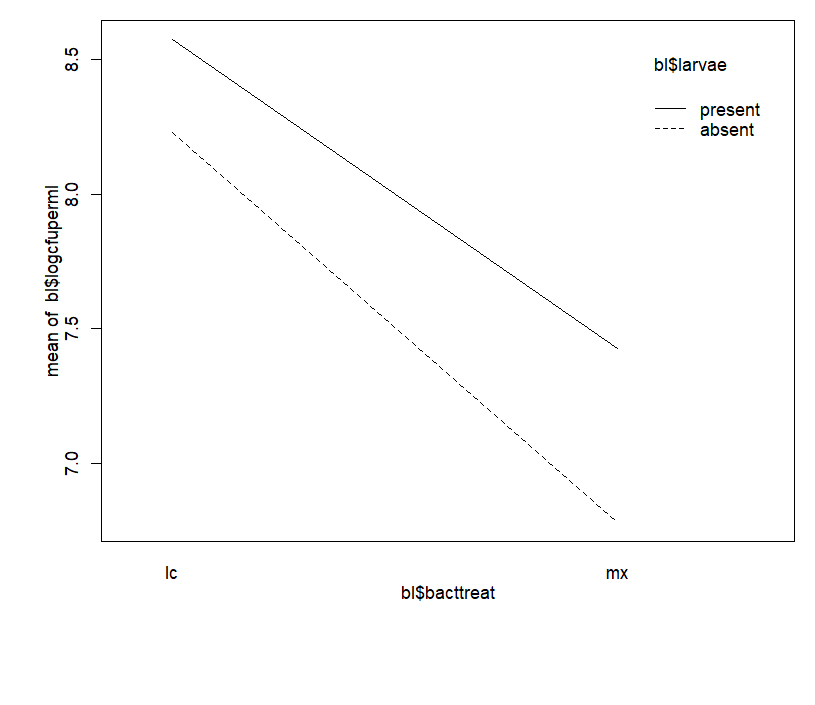

Mean log_10_(CFU per ml)

Microbiota treatment

Larvae

**Fig. S9.** Two-way interaction between bacterial treatment (MX vs. LC) and presence of larvae (absent vs. present) on bacterial load in larval rearing water

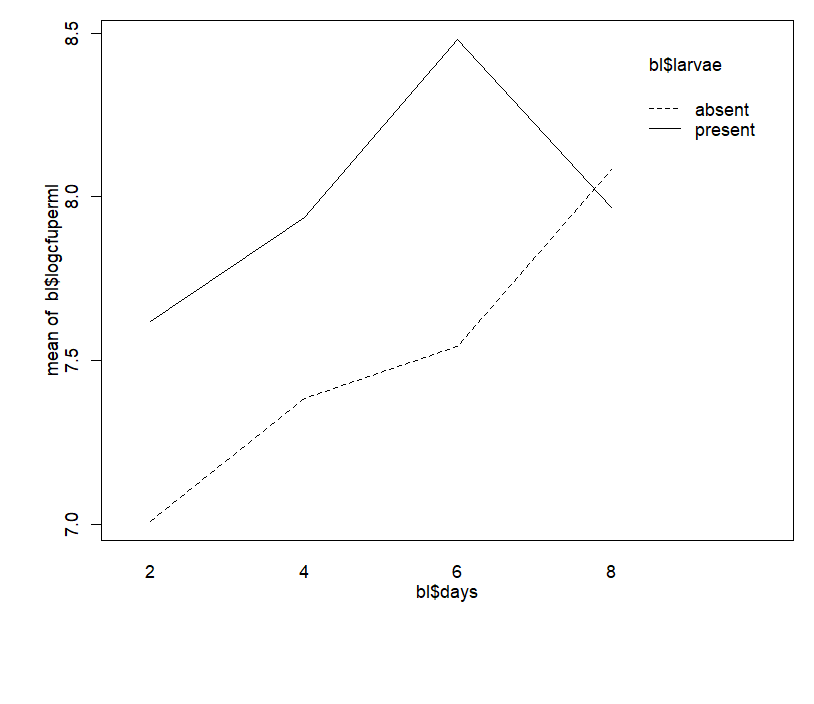

Mean log_10_(CFU per ml)

Larvae

Days

**Fig. S10.** Two-way interaction between presence of larvae (absent vs. present) and day on bacterial load in larval rearing water

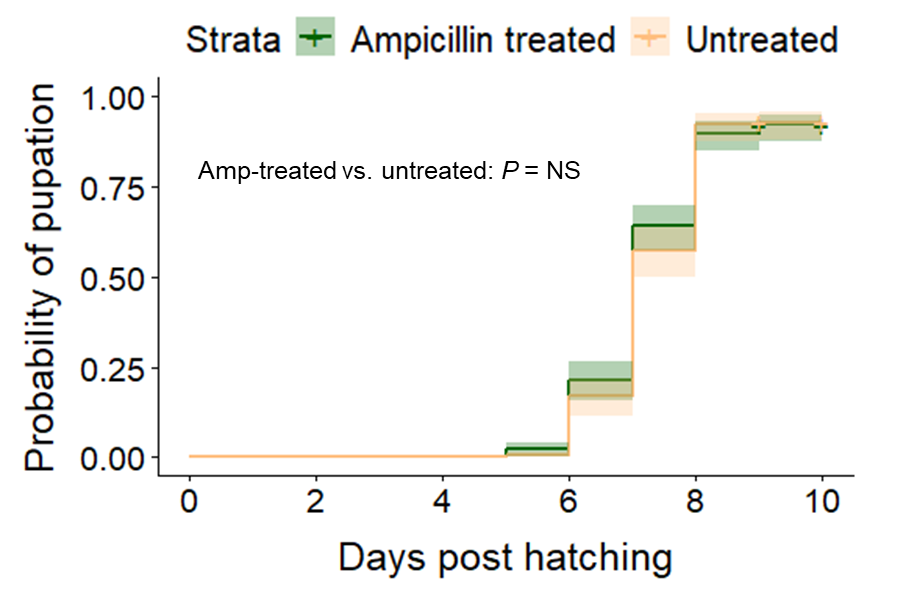

**Fig. S11.** Pupation rate was not affected by ampicillin treatment (*P* = 0.595). Pupation data were collected from three replicates, with *n* = 45–75 mosquitoes per treatment per replicate

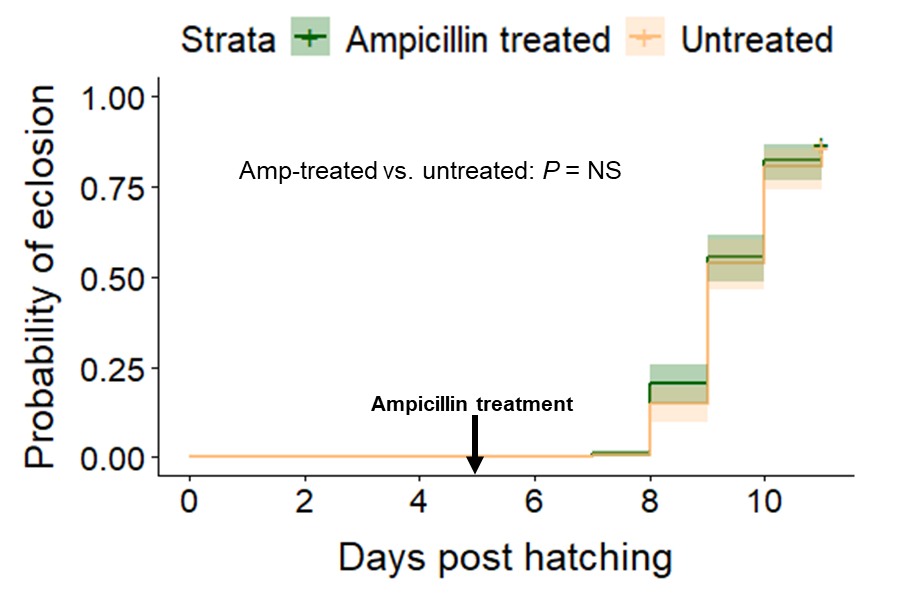

**Fig. S12.** Eclosion rate was unaffected by ampicillin treatment (*P* = 0.582). Eclosion data were collected from three replicates, with *n* = 45–75 mosquitoes per treatment per replicate
